# Cross-platform DNA motif discovery and benchmarking to explore binding specificities of poorly studied human transcription factors

**DOI:** 10.1101/2024.11.11.619379

**Authors:** Ilya E. Vorontsov, Ivan Kozin, Sergey Abramov, Alexandr Boytsov, Arttu Jolma, Mihai Albu, Giovanna Ambrosini, Katerina Faltejskova, Antoni J. Gralak, Nikita Gryzunov, Sachi Inukai, Semyon Kolmykov, Pavel Kravchenko, Judith F. Kribelbauer-Swietek, Kaitlin U. Laverty, Vladimir Nozdrin, Zain M. Patel, Dmitry Penzar, Marie-Luise Plescher, Sara E. Pour, Rozita Razavi, Ally W.H. Yang, Ivan Yevshin, Arsenii Zinkevich, Matthew T. Weirauch, Philipp Bucher, Bart Deplancke, Oriol Fornes, Jan Grau, Ivo Grosse, Fedor A. Kolpakov, The Codebook/GRECO-BIT Consortium, Vsevolod J. Makeev, Timothy R. Hughes, Ivan V. Kulakovskiy

## Abstract

A DNA sequence pattern, or “motif”, is an essential representation of DNA-binding specificity of a transcription factor (TF). Any particular motif model has potential flaws due to shortcomings of the underlying experimental data and computational motif discovery algorithm. As a part of the Codebook/GRECO-BIT initiative, here we evaluated at large scale the cross-platform recognition performance of positional weight matrices (PWMs), which remain popular motif models in many practical applications. We applied ten different DNA motif discovery tools to generate PWMs from the “Codebook” data comprised of 4,237 experiments from five different platforms profiling the DNA-binding specificity of 394 human proteins, focusing on understudied transcription factors of different structural families. For many of the proteins, there was no prior knowledge of a genuine motif. By benchmarking-supported human curation, we constructed an approved subset of experiments comprising about 30% of all experiments and 50% of tested TFs which displayed consistent motifs across platforms and replicates. We present the Codebook Motif Explorer (https://mex.autosome.org), a detailed online catalog of DNA motifs, including the top-ranked PWMs, and the underlying source and benchmarking data. We demonstrate that in the case of high-quality experimental data, most of the popular motif discovery tools detect valid motifs and generate PWMs, which perform well both on genomic and synthetic data. Yet, for each of the algorithms, there were problematic combinations of proteins and platforms, and the basic motif properties such as nucleotide composition and information content offered little help in detecting such pitfalls. By combining multiple PMWs in decision trees, we demonstrate how our setup can be readily adapted to train and test binding specificity models more complex than PWMs. Overall, our study provides a rich motif catalog as a solid baseline for advanced models and highlights the power of the multi-platform multi-tool approach for reliable mapping of DNA binding specificities.

Graphical Abstract

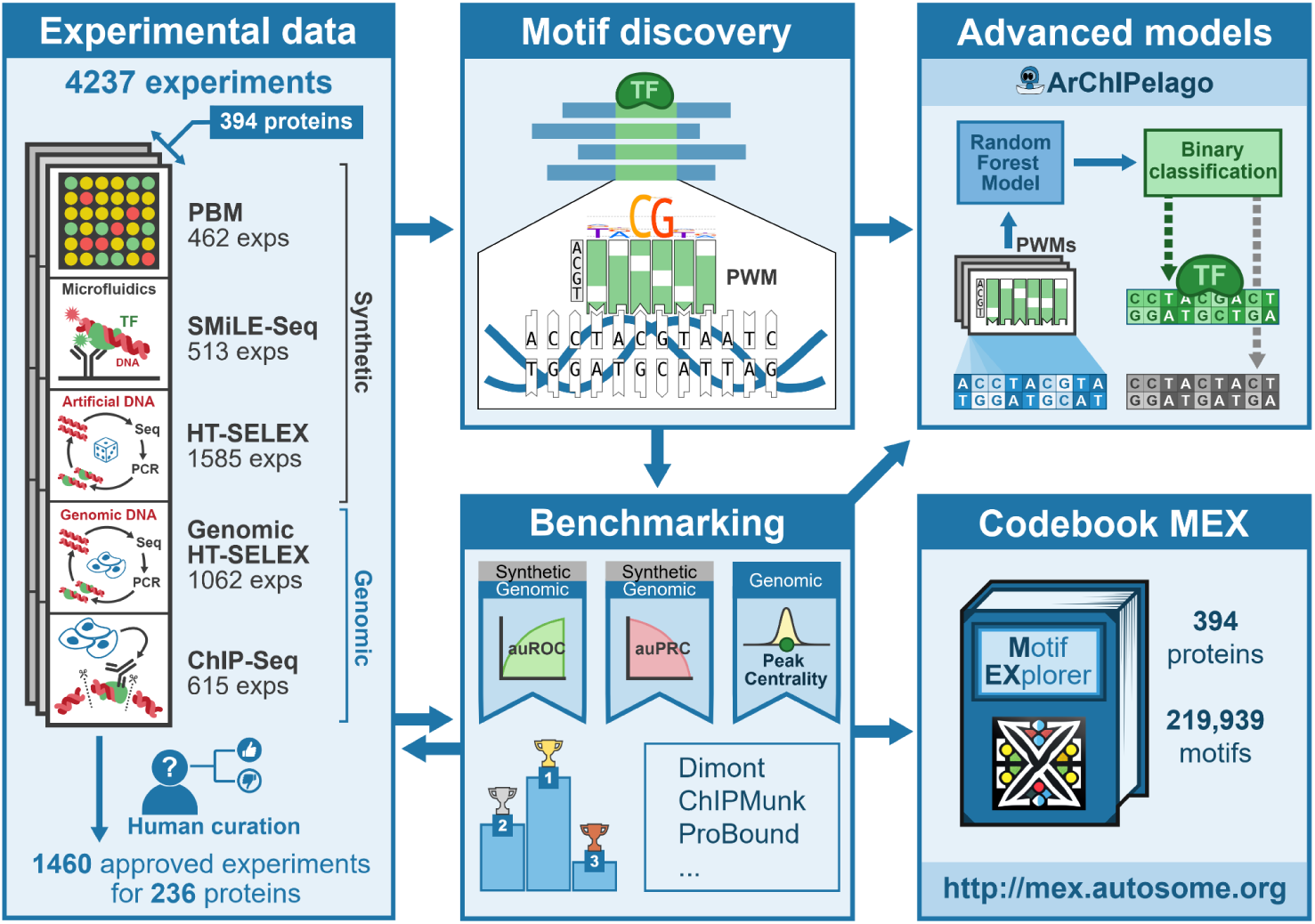

## Introduction

Transcription factor (TF) binding to DNA is a crucial component of transcriptional regulation, responsible for coordinated gene expression within gene regulatory networks^1^. Alteration of TF-DNA interactions is also a major cause of gene expression changes due to sequence variants in genome regulatory segments^2^. Thus, knowledge of DNA patterns, the “motifs”, specifically recognized by TFs, is essential for annotating gene regulatory regions^3^, interpreting regulatory variation^4^, and deciphering the logic learned by deep neural networks from genomics data^5^. The traditional and most widely used representation of a motif is the position weight matrix (PWM). It has few independent parameters (usually, 3 multiplied by the binding site length) and relies on a simple assumption: independent (additive) contributions of neighboring nucleotides to the overall binding energy^6^. Beyond PWMs, there are many advanced models of DNA sequences specifically recognized by TFs that account for interdependent nucleotide contributions^7^. Yet, despite the rapid progress of advanced machine learning applications in TF binding sites (TFBS) modeling, PWMs remain in the bioinformatics toolbox for studying gene regulation, and the databases providing TF binding motifs as PWMs, including CIS-BP^8^, JASPAR^9^, and HOCOMOCO^10^, are actively accessed and cited.

To construct a motif model such as a PWM, it is necessary to solve the motif discovery problem, i.e. perform pattern recognition in collections of related DNA sequences, such as those bound by TFs. A plethora of experimental approaches have been developed to identify TFBSs in random sequences, complete genomes, or their fragments^11^, and each of the approaches has its advantages and limitations. For example, the current high-throughput gold standard method for identifying TFBSs genome-wide *in vivo*, chromatin immunoprecipitation followed by sequencing (ChIP-Seq), has limited resolution and relies on computational analysis to delineate bound regions. While better resolution is achieved with ChIP-exo^12^ or CUT&RUN^13^, these methods still require motif modeling to pinpoint the exact binding site locations. Beyond specific interactions of a protein with particular nucleotide patterns, TF binding *in vivo* e.g., revealed by ChIP-Seq, is influenced by many cell type-specific factors: chromatin accessibility, availability of transcription cofactors, or competition with other proteins for the same binding regions^14^. Isolating the contribution of the DNA sequence from features of the cellular or genomic contexts is challenging, especially when considering genomic *in vivo* data only. Data obtained *in vitro* avoids these issues and, with synthetic sequences, it is possible to explore the sequence space more uniformly. However, these methods also have their own technical biases, for instance, high-throughput SELEX (HT-SELEX)^15^ saturates quickly with the strongest binding sequences^16^. Therefore, to overcome these challenges, the binding specificity of a TF ideally should be studied both *in vivo* and *in vitro* with both synthetic and genomic sequences, using multiple experimental platforms^17^.

Until now, there have been very few systematic studies evaluating the performance of different motif discovery tools on the outputs of different experimental assays. The well-known large-scale benchmarking of motif discovery algorithms conducted by Tompa *et al.* in 2005^18^ took place in the low-throughput era, and did not include any of the current experimental assays. A focused competition organized by Weirauch *et al.^19^* using *in vitro* protein-binding microarray (PBM) employed ChIP-seq as an external control but did not include motif discovery from experiments other than PBMs. Other studies either compared the performance of motif discovery tools only on simulated data^20^ or evaluated only pre-existing PWMs^16,17,21^.

Here we present the results of the Gene Regulation Consortium Benchmarking Initiative, GRECO-BIT, an offspring of the GRECO/GREEKC consortium^22^ dedicated to building and benchmarking algorithms for DNA motif discovery and TFBS modeling. Here, in collaboration with Codebook, we performed a large-scale motif analysis of newly generated human TF binding data^23^ obtained through five different experimental assays, using a variety of motif discovery tools followed by systematic benchmarking. Through comparative assessment of the resulting motifs, we developed the Codebook/GRECO-BIT Motif Explorer (Codebook MEX), an interactive catalog of motifs for 394 putative TFs that were analyzed in the Codebook dataset. This resource provides an overview of the efficacy of various tools for PWM-based motif discovery across different experimental platforms and highlights the PWMs with the highest overall rankings, thus laying the foundation for future benchmarking studies and paving the way for improved computational protocols for generating high-quality DNA sequence motifs.

## Results

In this study, we relied on the data from five experimental platforms used by the Codebook initiative^23^ to assay the binding specificity of 394 proteins (see ‘Workflow overview’ in **Methods**). The platforms included Chromatin immunoprecipitation followed by sequencing (ChIP-Seq^24^) and high-throughput SELEX with genomic DNA (GHT-SELEX^25^) to delineate genomic locations of TFBSs. The other three methods, standard high-throughput SELEX (HT-SELEX), selective microfluidics-based ligand enrichment followed by sequencing (SMiLE-Seq^26^), and protein binding microarray (PBM), were used to assess TF binding to synthetic DNA fragments with random sequences (e.g. 40N random inserts for HT-SELEX or pseudo-random probes in the case of PBMs^27^). For HT- and GHT-SELEX, there were three variants differing in the protein production method: GST-tagged *in vitro* transcription with *E. coli* extracts (-IVT), GFP-tagged IVT with wheat germ extracts (-GFPIVT), and whole human cell lysate (-Lys). Counting the variants of (G)HT-SELEX separately, at least two types of experiments were performed for 391 TFs. (see **Methods**, **Supplementary Table ST1,** and^23^). The experiments covered many previously unexplored or incompletely profiled TFs, thus complementing existing databases, as well as a number of well-studied TFs (positive controls). To our knowledge, this is the first time that such a large collection of TFs has been assessed in such a diverse set of experiments in parallel. This setup provides a unique opportunity for a cross-platform assessment of the performance of motif discovery tools with motifs derived from one experiment type tested with the data from other types of experiments.

For clarity, in this study, we use the term “motif” to refer to a DNA binding specificity pattern, and “PWM” to refer to a motif model with one of two interchangeable representations: (1) a matrix of normalized nucleotide frequencies (also called the Position Frequency Matrix, PFM) or (2) a log-odds position weight matrix^6^. We adopted a systematic approach for motif discovery and benchmarking (**Figure 1A**) starting with uniform preprocessing of the data, such as peak calling (for GHT-SELEX and ChIP-Seq data) and normalization (for PBMs), and splitting results of each experiment into training and test sets (see **Methods**). Then, in the first round of motif discovery, we applied nine software tools to the training data of all experiments. We used classic MEME^28^ software, popular bioinformatics tools from the era of high-throughput data (HOMER^29^, ChIPMunk^30^, Autoseed^31^, STREME^32^, and Dimont^33^), and advanced methods (ExplaiNN^34^ and, for selected datasets, RCade^35^ and gkmSVM^36^). Not all tools were compatible with all data types, e.g., RCade was exclusively used for zinc finger TFs, and a specialized adaptation of Dimont for HT-SELEX (DimontHTS) was used for HT-SELEX data.

**Figure 1.**
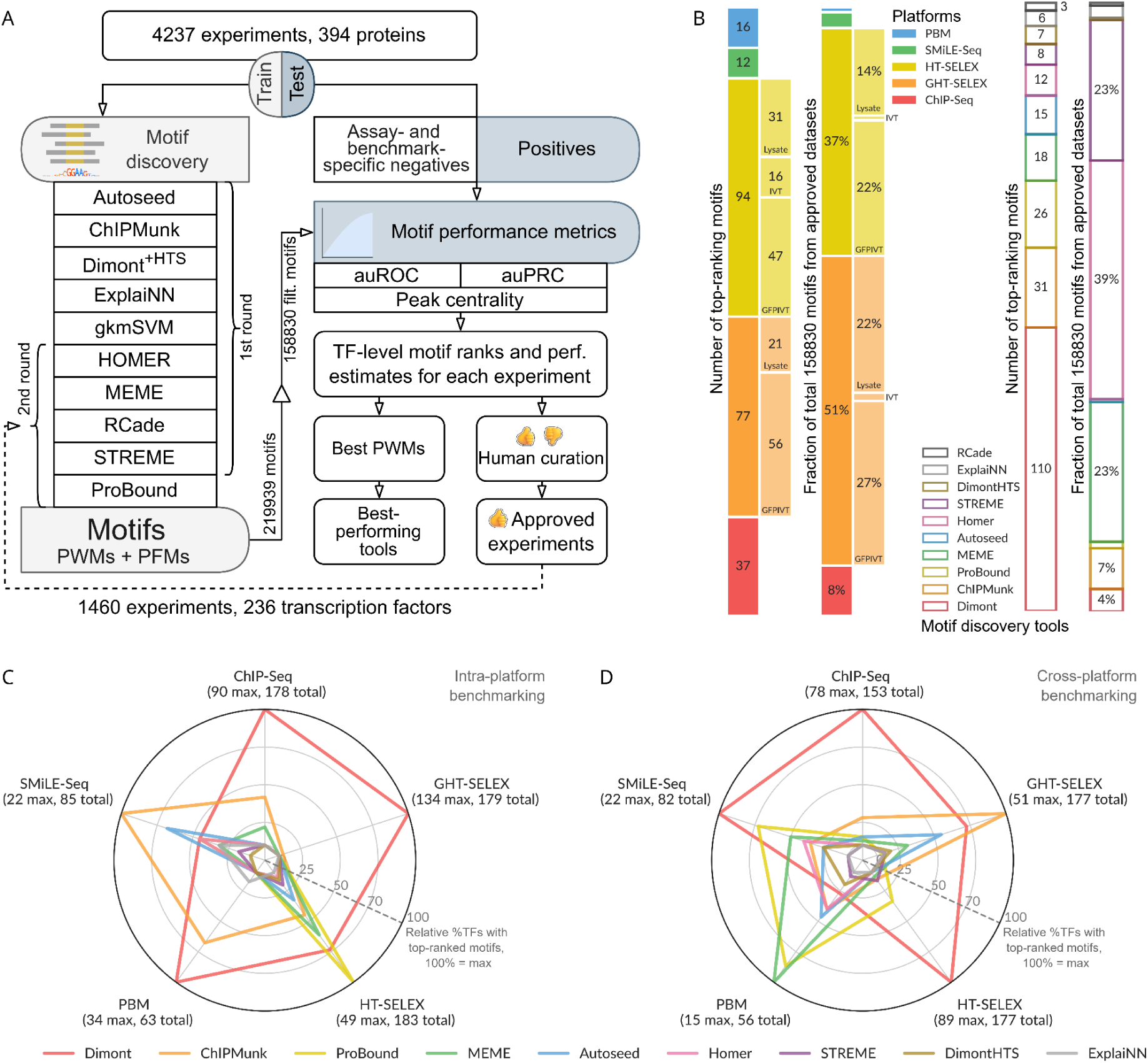
Motif discovery and benchmarking pipeline and the collection of top-ranking motifs. **A:** Schematic of the pipeline. **B:** Contributions of different tools and experimental methods (types of GHT-SELEX and HT-SELEX are shown in extra bars) to the top-ranking motif collection (numbers of TFs) and to the complete MEX set of benchmarked motifs (expressed as a percentage). **C, D:** Relative percentage of TFs with top-ranking motifs produced by a particular motif discovery tool compared to the highest number of top-ranking motifs yielded by a single tool (defines 100% at the radar axes). The maximum number of top-ranking motifs (one motif per TF) per tool for each data type is shown in parentheses, along with the total number of eligible TFs (with the train, test data, and motifs from the same platform). **C:** Intra-platform testing: the motifs are constructed and tested on the same type of experiment. **D:** Cross-platform testing, motifs obtained from all but one experiment type are tested on this experiment type of interest (labeled at the radar axes). As in panel C, only the top-ranking motifs for the selected TF are counted. TF: transcription factor, PWM: position weight matrix, PFM: position frequency matrix, auROC: area under the receiver operating characteristic, auPRC: area under the precision-recall curve.

With the diverse set of platforms and different motif applications, it is not trivial, if at all possible, to select a single universal benchmarking metric of motif performance. Thus, to evaluate the classification performance of all PWMs across the test data from all platforms, we employed multiple dockerized benchmarking protocols from^17^, with additions from^10^ and^37^, and adapted methodology for PBMs from^19^ (see **Methods**). Technically, to scan a DNA sequence with a given motif, most of the employed benchmarking protocols use the sum-occupancy scoring^16^. Specifically for ChIP-Seq and GHT-SELEX peaks, we also ran the HOCOMOCO benchmark^37^, which considers only a single top-scoring log-odds PWM hit in each sequence, and estimated the CentriMo motif centrality score, which accounts for the distance of the binding site to the peak summits^38^. To allow for streamlined applicability of existing benchmarking protocols, alternative motif representations from advanced methods were converted to PFMs and PWMs.

Many of the TFs were previously uncharacterized, and the initial benchmarking therefore served as the criterion to determine which experiments were successful. To this end, initial benchmarking results underwent human expert curation to approve a subset of successful experiments for detailed analysis. To approve the experiment, we required that either (1) motifs discovered from this experiment were consistently similar between platforms or similar to related known or Codebook TFs and scored highly in different benchmarks or (2) the motifs originating from and high-ranking on other approved experiments scored highly on the test dataset built from the experiment under curation. During curation (see **Methods**), we took into account known motifs both to validate real cases (positive controls and Codebook TFs from well-studied families) and to exclude artifacts, or “passenger” motifs" besetting multiple independent experiments. As a result, the approved set of experiments encompassed 236 TFs and comprised 1,460 datasets. To expand the motif sets from popular tools (HOMER, MEME, RCade, STREME) and explore another advanced method (ProBound^39^), we additionally ran a second round of motif discovery and benchmarking using the “approved” datasets.

In total, this effort generated 219,939 PWMs, with 164,350 derived from the approved datasets. Out of these, 158,830 PWMs passed additional automatic filtering for common artifact signals (such as simple repeats and the most widespread ChIP contaminants; see **Methods**) and formed the primary motif set for the downstream analysis (**Supplementary Figure SF1A, B**). The resulting resource, which includes the train-test sets and 16,812,803 performance estimates, 9,317,269 of which belong to the motifs from approved datasets, is available at ZENODO^40–42^. Interactive access to both the approved and complete motif collections, alongside the benchmarking results, is provided by the Codebook Motif Explorer (Codebook MEX, https://mex.autosome.org).

### Motif discovery from diverse experiments requires a diverse toolbox

Using the data from approved experiments, we sought to construct a global benchmarking ranking for different motif discovery algorithms across different experimental platforms. To this end, we employed a hierarchical ranking procedure to sequentially identify the top-ranking motifs for each TF and experiment type across multiple individual performance metrics, replicates, and types of experiments, followed by a global TF-level ranking (see **Methods**).

We expected that the benchmarking study would reveal different motif discovery tools to be either universally superior or best suited to derive motifs from particular types of experiments. However, these expectations were only partially met. Nearly all software tools contributed to the final collection of the globally top-ranking motifs per TF (**Figure 1B**), i.e., there was no algorithm, in retrospect, that was not worth including (The one possible exception was gkmSVM, which, due to high computational load, was applied only to a subset of ChIP-Seq data, resulting in the smallest initial set of motifs, and was tested here using derived PWMs, rather than its native sequence scanner). Nonetheless, nearly half of the top-ranking motifs in the final collection were generated by a single tool, Dimont. This dominance persisted even when considering the top 20 motifs for each TF (**Supplementary Figure SF1C**). Furthermore, the proportional contribution of the tools to the collection of top-ranking motifs did not reflect the initial quantity of motifs generated by these tools (**Figure 1B**). In contrast, the distribution of experiment types that yielded the top-ranking motifs was more similar to the original composition of the Codebook data: the top-ranking motifs were largely derived from ChIP-Seq and (G)HT-SELEX experiments (**Supplementary Table ST1**).

When considering individual experiment types, we first examined which tool produced the top-ranking motifs across TFs when trained and tested on data from the same type of experiment (**Figure 1C**, **Supplementary Figure SF2**). Dimont was the top performer in three categories (GHT-SELEX, ChIP-Seq, PBM), and was competitive for HT-SELEX, which constituted the majority of the data, thus explaining its significant contribution to the global set of top-scoring motifs. ProBound led in HT-SELEX, while ChIPMunk was the second best for many types of experiments, and led for SMiLE-Seq. This comparison highlights tools that excel at capturing experiment-specific motifs, potentially including biases and artifacts in addition to the intrinsic binding specificity of the TF.

Next, we conducted a cross-platform analysis, where we assessed TFBS prediction performance for a particular type of experiment using motifs discovered from all other types (**Figure 1D**, **Supplementary Figure SF2**). Considering top-ranking motifs, Dimont scored highest overall for SMiLE-Seq, HT-SELEX, and ChIP-Seq. Conversely, ChIPMunk excelled with genomic HT-SELEX and MEME with PBMs. Interestingly, ProBound was powerful in predicting PBM and SMiLE-Seq, and to a lesser extent, HT-SELEX, suggesting its ability to capture lower-affinity binding sites common in PBM and SMiLE-Seq data. Analyzing the subtypes of GHT-SELEX and HT-SELEX individually (**Supplementary Figure SF3**), the results were similar (Dimont led in GHT-SELEX and ProBound in HT-SELEX), and the observed variability between experiment subtypes likely reflects the differences in the profiled TFs and the signal-to-noise ratios of particular experiments.

Summing up, considering overall motif rankings, Dimont excels across the board and particularly at ChIP-Seq, but intra- and cross-platform benchmarking highlight alternative motif discovery tools best suited for TFBS recognition in specific scenarios.

#### The first motif reported is the best in benchmarking in 75% of cases

A single run of a particular motif discovery tool, be it MEME or Dimont, may yield multiple motifs, and ideally it should put the true binding motif at the top of the list. In some datasets, however, the first reported motifs may reflect the binding patterns of a TF cofactor (*e.g.* in ChIP-Seq) or even spurious signals such as artificially enriched sequences (*e.g.* aptamers in HT-SELEX). Yet, these examples are usually experiment-specific and unlikely to be ranking highly in the overall benchmarking across different platforms. Thus, motifs ranked higher in the overall benchmarking should have a higher probability of reflecting the true binding specificity.

For each run of a motif discovery tool on a particular dataset for a particular TF, we took the first three reported motifs (excluding common artifact signals, see **Methods**), and located them in the overall benchmarking ranking (that includes motifs from all programs and all datasets for the TF). In about 75% of cases (runs of a particular software using a particular training dataset that yielded more than one motif), in the global benchmarking, the first reported motifs indeed scored higher than the other two (**Supplementary Figure SF4A**). However, in the remaining 25% of cases, the highest-ranked motifs from a particular motif discovery run were not the first in the software output, i.e., the internal ranking of the motif discovery tools failed to distinguish what we assume are the proper signals. In real-life scenarios, this percentage could be even higher as *a priori* pre-filtering of common artifacts would not be possible without multi-platform data. Therefore, in practice, secondary motifs reported by motif discovery tools must be considered in downstream analyses. Of note, Dimont and ChIPMunk stood out in this test and for these tools, the first reported motifs were relevant in 90-95% of cases.

#### Quantitative analysis of motif performance

While ranking analysis provides a bird’s eye view, it does not reveal the actual performance difference between the winning and runner-up motifs. To compare the motifs and the motif discovery tools quantitatively, we introduced an overall metric of motif performance across different benchmarks and datasets yielded by a particular experimental platform. Briefly (see **Methods** for more details), for each quadruple of a TF, a dataset, a benchmark (e.g. binary classification of bound and non-bound sequences), and a performance metric (e.g. auROC) we rescaled the values across motifs into the range [0,1], 0 corresponding to the worst and 1 to the best of values achieved by different motifs. The overall performance of a motif is the average of those rescaled values across all metrics and experiments of a particular type. For each TF and each motif discovery tool, we then selected representative motifs achieving the highest overall performance in the intra- and cross-platform analysis with particular target experiment types.

**Figure 2** displays the median and the interquartile range (IQR) of the overall performance of representative motifs across TFs assayed in the intra- and cross-platform fashion (**Figure 2A** and **2B**). In these tests, we expected the tools with wider applicability across TFs to yield a higher median and lower IQR. In the intra-platform comparison, HT-SELEX appeared to be the most agnostic to the motif discovery tool, as the median overall performance was consistently high with low IQR across TFs for all tools. In contrast, the largest differences between tools were observed for SMiLE-Seq data. From the tool-centric perspective, the mean performance of Dimont, ChIPMunk, and Autoseed was stable across TFs and platforms, although MEME and HOMER were not far behind. RCade performance on SMiLE-Seq data is noteworthy, as it was applied only to a small subset of TFs, but the resulting motifs displayed a strong performance. Finally, ExplaiNN could not quantitatively compete with other tools when trained and validated on genomic regions, potentially due to a lower-than-necessary volume of available training data. In the cross-platform setting, the differences between the tools lessened: all conventional tools (MEME, STREME, HOMER, ChIPMunk, Dimont) performed comparably well. We rationalize this outcome by the fact that the representative motifs were collected across platforms allowing each tool to avoid individual within-platform pitfalls, although gkmSVM displayed acceptable median results despite being trained on ChIP-Seq data only. Still, the IQR was high for many specific combinations of tools and experiment types, i.e., depending on a TF, any single tool can fail to properly capture a universally applicable motif even with multiple attempts across platforms, despite performing satisfactorily on average.

**Figure 2.**
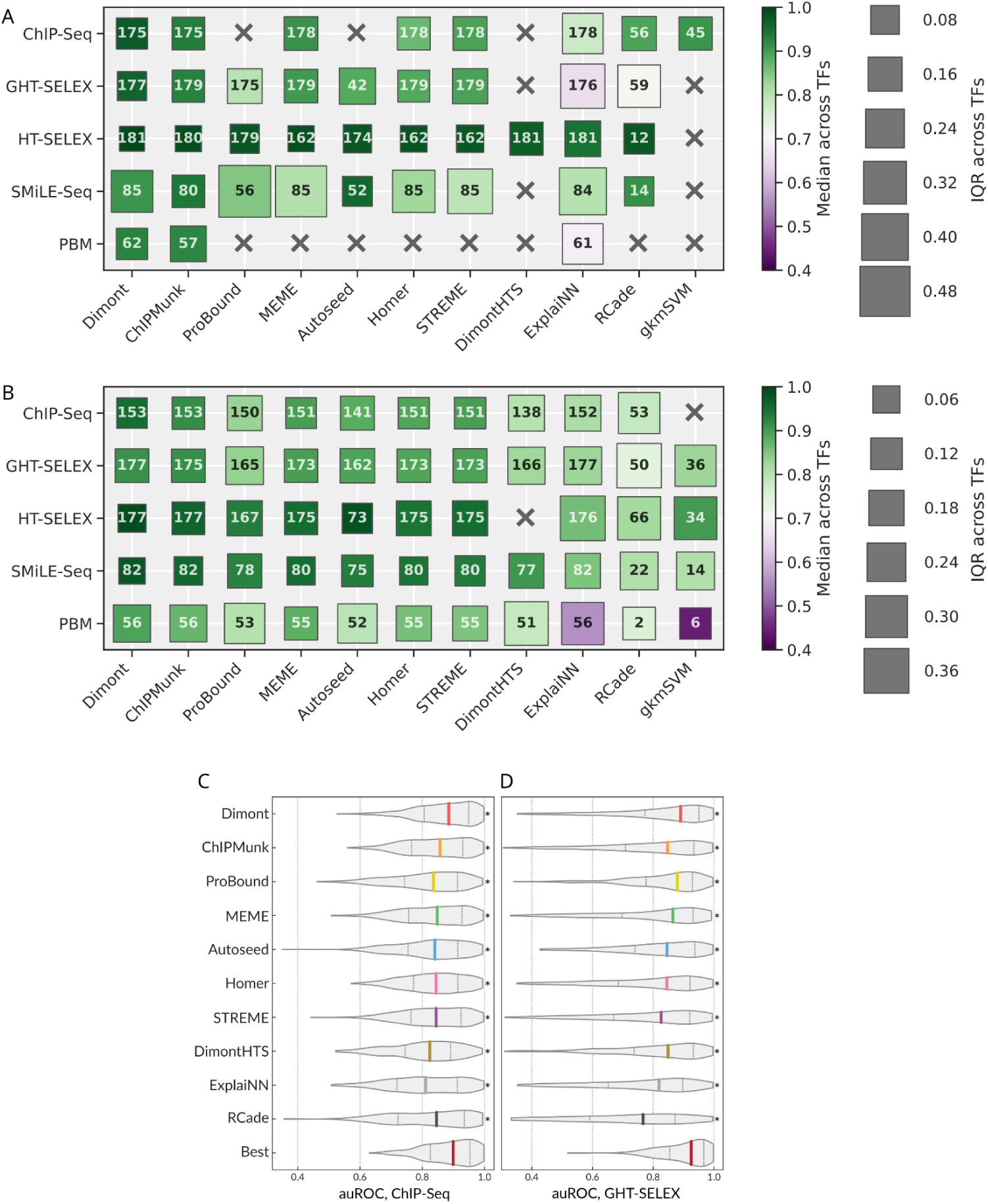
Quantitative analysis of the intra- and cross-platform performance of different motif discovery tools. **A:** Highest overall performance of the best motifs (one per TF) when training and testing on the same type of experiment. **B:** Highest overall performance of the best motifs (one per TF) in cross-platform evaluation. The color scale (identical in A and B) represents the median performance (higher is better), and the size of the boxes (note the different scale between A and B) indicates the IQR (lower is better) across TFs. The number in each square shows the total number of tested TFs for each combination of a motif discovery tool and an experiment type. **C, D:** Distributions of auROC values for all TF-dataset pairs calculated from the top-ranking motifs from each motif discovery tool selected by global benchmarking: tested on ChIP-Seq (C), tested on all variants of genomic HT-SELEX (D). The bottom violin is built from the highest values obtained for different TF-dataset pairs considering the top-ranking motifs from all tools. auROC: area under the receiver operating characteristic, IQR: interquartile range, TF: transcription factor.

The scaling used to obtain the overall motif performance estimates conceals the information on the absolute efficacy of a tool. Indeed, even if all the tested motifs are of very high quality and achieve the auROC over 0.9 on a particular dataset, the respective scaled values are still stretched across the [0,1] band. To obtain more interpretable performance estimates, we plotted the distribution of the raw values of the area under the receiver operating characteristic (auROC, computed with sum-occupancy PFM scoring as in^17^) using top-ranking motifs from each tool. Violin plots illustrate the distribution of raw auROC values across TFs and test datasets for ChIP-Seq (**Figure 2C**) or GHT-SELEX (**Figure 2D**) data. The values reached by Dimont are consistently higher than those of other tools and close to those of the best PWMs (selected irrespectively of the tool), with the median auROC across TFs and datasets over 0.85 for both ChIP-Seq and GHT-SELEX. A different performance metric, asymptotic pseudo-auROC (computed with best log-odds PWM best hits as in^37^), was less discriminative between tools (see **Methods** and **Supplementary Figure SF4B**).

The diversity of experimental platforms allows for answering another important question: whether protein binding experiments with synthetic oligonucleotides provide motifs suitable for reliable prediction of genomic binding sites. For different TFs, and different datasets, we plotted the distributions of difference between the maximal auROC values achieved with the genomic data (ChIP-Seq and GHT-SELEX) by motifs obtained from the genomic and the synthetic data (HT-SELEX, SMiLE-Seq, PBM), **Figure 3A, B**. Overall, we observed a visible drop of auROC median of -0.1 to -0.2 depending on the tool, meaning that the genomic binding sites remain difficult to predict with motifs from synthetic data even when using multiple motif discovery tools. In extreme cases, for some datasets, the auROC dropped extremely low (ΔauROC < -0.5). However, there were many cases with only a marginal decrease of auROC as the area around ΔauROC near 0.0 is densely populated for many motif discovery tools.

**Figure 3.**
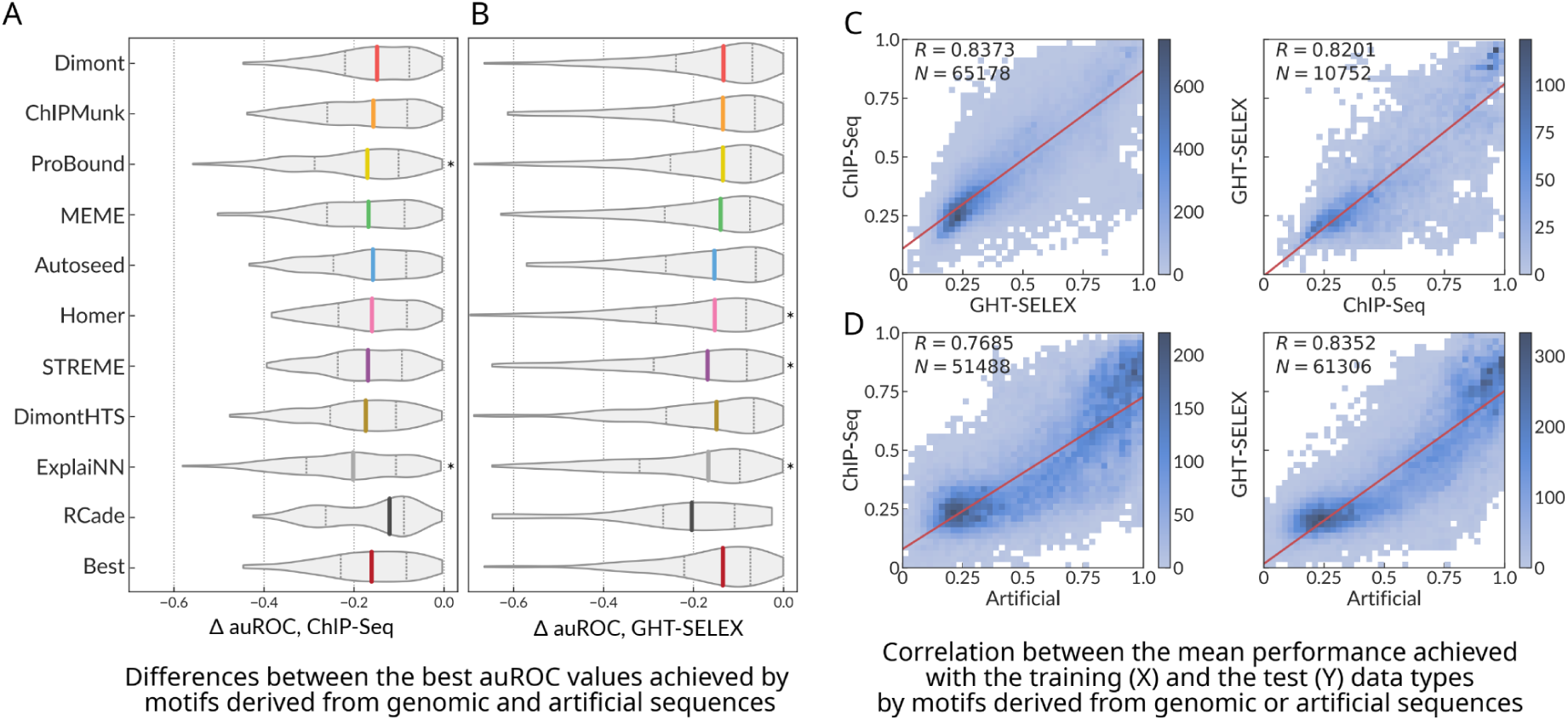
Performance of motifs derived from artificial sequences when applied to prediction of genomic binding sites. **A, B**: The difference between the best auROC values for genomic data achieved by motifs discovered in the genomic (ChIP-Seq or GHT-SELEX) or artificial (HT-SELEX, SMiLE-Seq, PBM) sequences; ChIP-Seq (A), GHT-SELEX (B). The last violin at the bottom shows the difference between the highest best-achieved values for each TF across all tools. *p < 0.05, paired Wilcoxon test against the best-achieved values (the bottom violin). IQR: interquartile range. **C, D:** Correlation between the overall performance of motifs at the training data (ChIP-Seq, GHT-SELEX or any artificial, X axes) and the test data (ChIP-Seq or GHT-SELEX, Y axes).

A thorough analysis of the overall performance of motifs for all pairs of platforms (**Supplementary Figure SF5**) reveals that there are subsets of motifs achieving very high scores at the training and the test experiment types simultaneously for almost any “training-test” combination of platforms. In particular, there are motifs identified from the synthetic data, which perform equally well on the genomic data (**Figure 3C, D**). Further, the overall performance at the training and the test data types are highly correlated, suggesting that motif performance measurements with successful experiments from any platform are predictive regarding motif applicability to other data and that high-scoring motifs likely provide a good and generalized representation of the true binding specificity of a TF of interest. We caution, however, that there are multiple outliers; thus, high performance for a single training data type does not absolutely guarantee universal cross-platform transferability.

Considering individual tools, their performance generally followed the global trend, except for an unexpectedly good RCade performance when tested on ChIP-Seq but not GHT-SELEX data (**Figure 2C**, **Figure 3A** versus **Figure 2D**, **Figure 3B**). This effect permits a simple explanation: by design, RCade obtained motifs for zinc-finger TFs only. For these TFs the performance ratings at the respective genomic datasets were higher for many other tools (**Supplementary Figure SF4C**), and binding sites in ChIP-Seq datasets were easier to predict than those in GHT-SELEX (**Supplementary Figure SF4D**).

#### Interpreting the role of flanking regions with motifs from gkmSVM

GkmSVM was used in the first round of motif discovery from ChIP-Seq data, covering 45 TFs with approved datasets. As gkmSVM was computationally demanding and its motifs were not top-ranking, we did not apply it to the analysis of the entire Codebook collection. However, gkmSVM motifs trained on the ChIP-Seq data performed competitively (**Figure 2A**). By examining the motifs of the 45 TFs constructed with GkmExplain from gkmSVM results, we found that it captured long sequence contexts in the vicinity of the binding sites, as seen from the motif length distribution (**Supplementary Figure SF6A**). Given the good performance of gkmSVM on ChIP-Seq and its acceptable performance for GHT-SELEX (**Figure 2B**), we concluded that the extended genomic context provided added value, at least for some TFs. Yet, the longer genomic context of binding sites is likely to represent properties of regulatory regions at a larger scale, including binding sites of other interacting TFs, rather than the genuine binding specificity of the protein under study.

#### Basic motif features are not related to motif scores in benchmarking

Basic motif features like motif length, information content (IC), and GC composition were irregular even for motifs derived with the same tool or experiment type. For instance, Dimont and ProBound produced many low-information content motifs, and for many tools, there were discrete spikes in preferred motif lengths arising from technical parameters or other technicalities of the motif discovery procedure (**Supplementary Figure SF6A, B**). The performance metrics were only weakly correlated to these basic motif features, however, as previously observed^17^. Considering genomic data, performance metrics were not correlated with GC composition or positional IC but did show a weak correlation with the motif length (Pearson ρ of 0.05 to 0.2, **Supplementary Figure SF7A**). This finding could reflect an ability of longer motifs to partially account for the contribution of the genomic context, as motif length was especially beneficial in the benchmark that used only the single best PWM hit per sequence (pseudo-auROC). In this scenario, an extra motif length optimization step could improve the resulting PWM performance for some tools^43^, although there is some risk that the motif does not represent the intrinsic activity of the TF: in benchmarks on artificial sequences, there was no correlation between motif length and performance at all, i.e., longer motifs did not have any advantages (**Supplementary Figure SF7B**) and only a very weak correlation (ρ around 0.06) with the positional IC. Focusing on the latter, a large IC spread was found even considering only the top 10 motifs for each TF (**Supplementary Figure SF7C**). Moreover, in our cross-platform assessment, the information content was not related to motif performance and instead reflected the motif origin experiment and motif discovery algorithm (**Supplementary Figure SF6**), suggesting that the average information content of a given high-scoring motif depends not only on the experimental platform and specifics of a particular experiment (e.g. sequencing depth, TF concentration, or signal-to-noise ratio) but also on the technical procedure employed for motif discovery. The biophysical explanation for the success of many low IC motifs is uncertain, but one hypothesis that has been put forward is that they average multiple binding modes^44^.

### A random forest of PWMs improves prediction of genomic binding sites

Many TFs display several modes of DNA recognition with clearly distinguishable motif subtypes^10,11^, and a straightforward advanced model could account for alternative TF binding modes by applying a logistic regression^45^ or decision trees^46^ on top of predictions from a collection of related PWMs. The richness of data provided by Codebook allows for a deeper exploration of such strategies.

Here we exploited the high similarity of GHT-SELEX and ChIP-Seq data and considered the genomic TFBS prediction task for 140 TFs for which both GHT-SELEX and ChIP-Seq experiments were approved and generated enough peaks for analysis (see **Methods**). As evaluating advanced models is more sensitive to biases in the data, we created dedicated train-test datasets with several negative controls, including “shades” (*i.e.* peak-neighboring regions as in the main PWM benchmarking), “alien” peaks of non-relevant TFs, and “random” genomic regions. The latter two background sets were built by sampling from available sequences to achieve the same distribution of the GC composition as that of the positive set. Using the random negative set, we trained the Archipelago model (see **Methods**), a random forest classifier built on top of best hits of a relatively small collection of log-odds PWMs (excluding those discovered from the test data type). Next, we estimated its mean performance with the three alternative negative sets and used PWMs reaching either the highest auROC or auPRC as the respective baselines.

#### Evaluation within experiment type

To avoid information leakage, each time, we excluded PWMs obtained from the test experiment type. Yet, it was still possible to train and test Archipelago with the single experiment type using PWMs from other platforms. In this scenario, Archipelago consistently outperformed individual PWMs, with significant gains in both auROC and auPRC across TFs (**Supplementary Figure SF8A-F**, with the average of ΔauROC and ΔauPRC shown in **Figure 4A**), although the random forest model showed only a marginal improvement over logistic regression (**Supplementary Figure SF8G**). In agreement with the primary PWM benchmark, the absolute intra-platform auROC values were high even for individual PWMs (**Supplementary Figure SF8A, D**), while achieving high auPRC values was more difficult due to class imbalance. As expected, the negative set made of shades, which had a less skewed class imbalance, has yielded the highest auPRC scores (**Supplementary Figure SF8B, E**). Overall, comparing different TFs, the improvement of Archipelago over PWMs did not depend on the size of the positive set or the number of PWMs in the model. Remarkably, only 2-4 PWMs combined were already sufficient for a major quality boost over a single PWM (**Supplementary Figure SF8H, I**).

**Figure 4.**
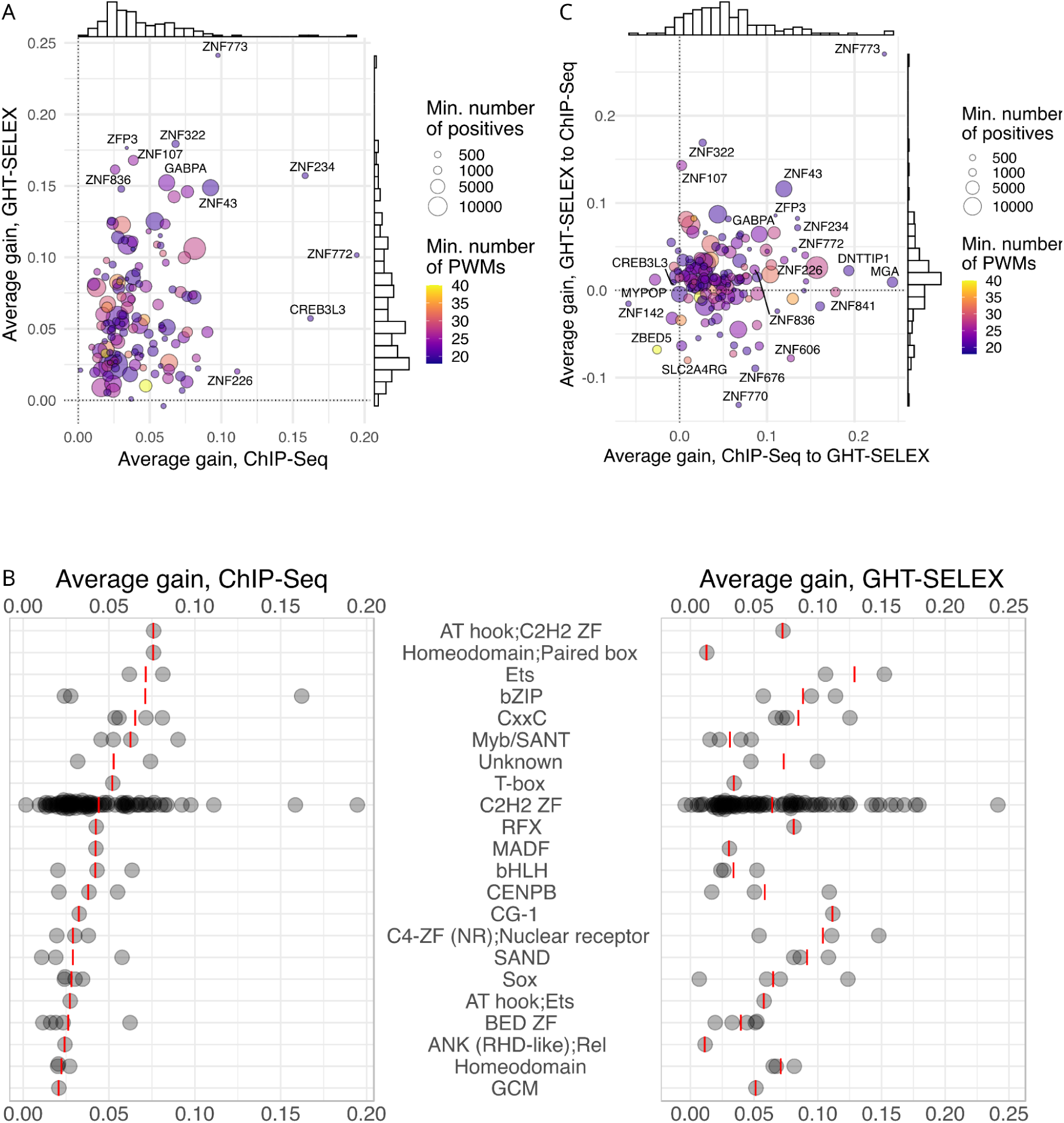
Improved prediction of binding sites with a Random Forest of alternative motifs (Archipelago). **A:** Average gain (ΔauPRC and ΔauROC averaged) between the best PWM and Archipelago. X-axis: training and test with ChIP-Seq, Y-axis: training and test with GHT-SELEX, color scale: the number of PWMs included in the model (minimum of GHT-SELEX and ChIP-Seq), point size: the size of the training positive set (minimum of GHT-SELEX and ChIP-Seq). **B:** Average gain achieved for different TF families. Red lines denote mean values, families are sorted by the median gain for ChIP-Seq. **C:** Average of ΔauPRC and ΔauROC between the best PWM and Archipelago when training and testing on different experiment types. X-axis: transfer from ChIP-Seq to GHT-SELEX, Y-axis: transfer from GHT-SELEX to ChIP-Seq.

The average performance gain for GHT-SELEX and ChIP-Seq differed for individual TFs and TF families (**Figure 4B**). Some TFs, such as GABPA, which is known to form multimers on DNA^47^, showed improvement with Archipelago presumably due to multiple distinct binding sites per peak not being captured by a single best PWM hit. Other TFs with clear improvement had low-complexity or low-information content motifs (*e.g.* ZNF772 and ZNF773), likely reflecting complex patterns that are hard to represent by a single fixed-width PWM. A comparable improvement was observed for several PWMs representing overlapping patterns (*e.g.* for CREB3L3), which may represent different half-site patterns; the improvement on ChIP-Seq may also fortuitously capture heteromeric sites^48^. Finally, some TFs bind several site subtypes, such as C2H2 zinc finger TFs with modular binding specificities^25^. A strong example is ZNF43 with PWMs representing single and double-box binding motifs which are conveniently taken into account together by Archipelago but not by single PWMs.

#### Evaluating the models’ transferability

At first glance, the cross-platform performance of Archipelago (training the RF on the ChIP-Seq and testing on GHT-SELEX and *vice versa*) seems contradictory: ChIP-Seq-to-GHT-SELEX train-test yields a stable performance increase, while the train-test in the opposite direction behaves more randomly and often underperforms even in comparison to the best single PWM (**Supplementary Figure SF9A-H**, **Figure 4C, Supplementary Table ST2**). This can be explained by comparing the training data volume available from ChIP-Seq and GHT-SELEX, with the latter generally providing 2-3 times fewer peaks (**Supplementary Figure SF9I**). This difference not only made the model training more prone to overfitting but also provided less information on the actual diversity of genomic binding sites. Another issue arises from different motif subtypes preferably represented in ChIP-Seq and GHT-SELEX peaks, e.g., for ZNF770, a longer motif variant is present in ChIP-Seq but is less prominent in GHT-SELEX. Yet, in the end, the top TFs receiving the highest performance gain in the cross-platform evaluation were partly shared with those with the highest performance gain in the intra-platform setup, such as ZNF773, ZNF43, and GABPA.

### Codebook Motif Explorer

The results of this study are presented through the interactive Codebook/GRECO-BIT Motif Explorer (MEX, https://mex.autosome.org), which provides motifs, performance metrics, ranks, logos, and structured downloads, such as sets of top-performing motifs and related metadata, see an overview in **Supplementary Figure SF10**. The complete set of MEX motifs and the benchmarking-ready Codebook data are also available at ZENODO^40–42^.

## Discussion

Computational methods for motif discovery in DNA sequences have been evolving for more than three decades, stimulated by progress in experimental methods for profiling DNA-protein interactions. Yet, quantitative assessment of the performance and reliability of motif discovery tools is lagging, partly due to a shortage of uniform data sets for validation between experimental platforms. Thus, forty years after the inception of PWMs and following many advances in the measurement and representation of DNA sequence specificities, it remains controversial how to best measure, derive, use, and test motif models. Further, there was no commonly accepted set of PWMs that could serve as a reliable baseline. This deficit complicates a fair assessment of alternative PWMs or comparison of PWMs to more complex models, able to account for correlations of nucleotides within binding sites, even for widely studied TFs with well-described DNA binding specificities. Particularly, in some applications the complex models can fall behind carefully selected PWMs^49^, but, without the commonly accepted baseline, it remains problematic to provide reliable quantitative estimates of the models’ added value.

In our study, we used multiple motif discovery tools and multiple performance metrics across multiple experimental platforms. We identified motifs that were ranked consistently high by all metrics and across multiple replicates and individual test datasets, such as SELEX cycles or test data from alternative PBM normalization strategies. This approach allowed us to avoid prioritizing motifs that received high scores in a single benchmark by chance: different performance metrics were positively correlated but agreed imperfectly and produced different motif rankings (**Supplementary Figure SF11**). Thus, we consider that the Codebook MEX motif set and the underlying data provide a valuable resource for further development of DNA motif discovery tools. Of note, in our study, we employed several advanced motif discovery methods, including gkmSVM, ProBound, and ExplaiNN. However, while testing them against classical tools, we reduced their efficacy by converting their results to simple PWMs. Therefore, it remains of interest to perform a dedicated study of advanced motif models using the MEX PWMs as a baseline.

A related problem arises with human curation of the experimental datasets (which in our case was essential) being based on PWM-represented motifs: there might be TFs with intricate DNA-binding specificity that cannot be captured by PWMs, making it impossible to properly assess and approve the dataset. Finally, by design, we did not balance the starting motif sets across tools, allowing multiple candidate motifs to enter the benchmarking pipeline. Although running a conventional tool multiple times and collecting multiple alternative motifs might be a common practical scenario when performing an exploratory analysis of TF-DNA binding specificity, this is not a fully suitable approach in terms of benchmarking. Yet, in our study, larger motif sets did not enable the respective tools to occupy the podium despite having ‘more attempts’ (**Figure 1B-D**, **Figure 2A, B, Supplementary Figure SF1**).

Dimont was the most versatile tool for PWM motif discovery, achieving the best performance on the entire experimental data set, while ChIPMunk and ProBound could, in some cases, compete with Dimont for ChIP-Seq/GHT-SELEX and HT-SELEX. Classical tools such as MEME and HOMER rarely gave the best-performing motifs but often provided stably good results, as the gap in absolute performance from the best motifs was neither borderline nor dramatic. The most notable difference of Dimont compared with most alternative motif discovery approaches is that the PWM model in Dimont is optimized for a discriminative objective function numerically instead of using count-based statistics. Depending on the experimental method, ‘discriminative’ may refer to distinguishing bound from unbound sequences, or to reconstructing a continuous scale of signals related to binding strength. Hence, Dimont optimizes its PWM using an objective function - though on the training data - that is in some sense related to the performance metrics that have been used for evaluation on the test data in the benchmark. Of note, while ChIPMunk, Dimont, ProBound, and ExplaiNN were applied by their authors, other tools including MEME and HOMER were employed not from their creators’ hands, and thus might be technically handicapped.

It is important to count failures as well as successes. While most of the tools performed well on average, particular combinations of TFs, tools, and types of experiments could be more problematic, and such cases are difficult if at all possible to detect *ab initio*. Importantly, our study benefited greatly from multiple types of experiments for a single TF, which is quite rare in practice, where researchers usually limit themselves to a single assay. In real-life scenarios, the success rate of motif discovery will be even lower and simultaneously harder to assess, as in our study we discarded more than half of the experiments as “non-approved”, and human curation would have been much more error-prone without the availability of data from multiple experimental platforms.

Our large-scale motif discovery and benchmarking efforts also highlighted the widely debated topic of whether innate TF binding specificity or genomic context (e.g., DNA bases flanking the directly contacted binding site) contribute more to the genomic binding profile of a TF. In our observations, the motifs learned from artificial sequences were highly transferable to genomic regions, thus reinforcing the idea that innate DNA binding specificity is localized, highly sequence dependent, and can be efficiently learned and modeled from diverse experimental data including synthetic DNA sequences. The property of many TFs to recognize several motif subtypes can be addressed by “ensembling” PWMs in the Random Forest classifier, which was able to achieve better performance when trained and tested on GHT-SELEX and ChIP-Seq peaks, suggesting that cell type-agnostic, reliable binding profiles can be generated *in silico* from DNA sequence alone^25^. However, it was significantly more difficult for models to generalize beyond the particular experimental platform, even though the underlying motifs were built from diverse experiments. Thus, we expect transferability and generalization to remain major challenges for machine learning applications in advanced TFBS modeling, and their success will likely depend on creative approaches for data integration from multiple experiment types.

Despite the richness of the underlying data and multiple benchmarking protocols involved, we caution against overestimating the potential of the PWM as a model. First of all, many TFs, especially those with multiple zinc fingers, or several DNA binding domains of different classes, can recognize alternative motifs or motif subtypes, which are impossible to capture with a single PWM model. Thus, with PWM-based motif discovery, some motif subtypes can be mixed or missing, and the benchmarking results might be biased towards either primary subtypes or mixtures of alternative patterns. Next, the scoring method is important: as in^17^, most of our benchmarking protocols used the sum-occupancy scoring^16^, which effectively accounted for the contribution of multiple binding sites. The ChIP-Seq and GHT-SELEX benchmarks with the best log-odds PWM hits were less sensitive, resulting in smaller differences between tools (**Supplementary Figure SF4**), particularly, Dimont lost its edge. In this sense, selecting the best motif discovery tool is less crucial if the goal is to pinpoint the single best binding site within a known binding region.

Of note, only 236 of the 394 putative TF proteins examined in our study yielded approved datasets. We explored multiple motif discovery tools but used a deliberately conservative human curation protocol. Thus, we encourage the community to further explore the remaining data with advanced models or more sophisticated preprocessing strategies, as it should be possible to successfully discover motifs from some of the non-approved datasets, as has been shown for SMiLE-Seq^26^.

The most recent attempt to rigorously catalog human DNA-binding TFs^50^ assigned the respective GO term to 1,435 human proteins, requiring strict experimental evidence of both the role played by the particular protein in transcriptional regulation and its DNA-binding specificity. The set of motifs curated and rigorously analyzed in our study provides strong evidence for DNA-binding specificity for 54 additional proteins, which should now become prime candidates for genuine sequence-specific DNA-binding human TFs.

## Methods

### Workflow overview

The study relies on results of diverse experiments performed by Codebook to assess DNA binding specificity of human transcription factors: ChIP-Seq (CHS), HT-SELEX with random DNA (HTS), HT-SELEX with genomic DNA (genomic HT-SELEX, GHTS), protein-binding microarrays (PBMs), and SMiLE-Seq (SMS), see^23^. For PBMs, two results from two alternative designs (ME and HK) were available. For HTS and GHTS, there were three distinct experimental designs, which were differing in target protein production, namely, *in vitro* transcription (-IVT), GFP-tagged IVT (-GFPIVT), and cell lysate (-Lysate). For SMiLE-Seq, in addition to the new Codebook experiments, we included 27 previously published SMS datasets as additional positive controls^39^. In comparison to the main Codebook study^23^, we additionally used new SMiLE-Seq data for TFE3 (positive control) and ZNF346 (SMiLE-Seq negative control, RNA-binding protein with non-specific DNA binding), explicitly considered the data for GFP (negative control), and excluded BAZ2A and REXO4 at the earlier stage as they did not yield sufficient sets of ChIP-Seq or GHT-SELEX peaks and their other experiments were also deemed unsuccessful. Some ChIP-Seq and GHT-SELEX experiments were deemed non-approved prior to benchmarking and curation as they did not yield a sufficient number of peaks: we required that the experiment must yield 50 or more technically reproducible peaks at the even-numbered autosomes used for motif benchmarking, see below. Also, we considered technical sequencing replicates independently as they yielded overlapping but not identical peak sets. The complete starting set contained results from 4237 experiments (including technical sequencing replicates) for 394 proteins, see **Supplementary Table ST1**.

The general workflow of the study is shown in **Figure 1A**. Briefly, the experimental data were preprocessed, split into training and validation data, and passed to the first round of motif discovery with nine different software tools (see details below). While all tested tools were generating PWMs in the end, they represented two distinct categories. (1) Classic probabilistic and enumerative motif discovery tools: Autoseed^31^, ChIPMunk^30^, HOMER^29^, MEME^28^, and STREME^32^. (2) Advanced tools utilizing probabilistic discriminative learning (Dimont^33^), protein sequence information (RCade^35^), and modern machine learning techniques (ExplaiNN^34^, gkmSVM^36^ followed by GkmExplain^51^). For motif benchmarking, depending on the particular protocol (see below), we used sum-occupancy scoring with PFMs or best hits of log-odds PWMs.

Results of the 4237 experiments for 394 proteins were used in the first round of motif discovery with nine tools. The motifs were benchmarked and their logos, along with quantitative performance metrics for recognizing binding sites across multiple datasets, were used for expert curation of the datasets, see the details below. The 1460 curated and approved datasets for 236 TFs were then used for the second additional round of motif discovery with the addition of ProBound^39^ and extra motifs generated by the conventional tools (MEME, HOMER, RCade, STREME) using alternative settings, see below. Motifs from the second round were also benchmarked and put together with the results from the first round for the curation-approved datasets. Of note, motifs highly similar to common experimental artifacts were filtered before benchmarking (see below). The software implementation of the data processing pipeline is available on GitHub (https://github.com/autosome-ru/greco-bit-data-processing).

The Codebook Motif Explorer (MEX) website (https://mex.autosome.org) provides motifs from both rounds of motif discovery and all datasets, while only motifs originating from the curation-approved datasets were included in the comparison of tools and experimental methods presented in this study. MEX motif logos were drawn with drawlogo (https://pypi.org/project/drawlogo/), which uses discrete information content^30^ and inflated pseudocounts to achieve visual clarity for low-information content motifs.

### Experimental data preprocessing

An overview of the experimental data and preprocessing pipeline followed by motif discovery and benchmarking is shown in **Figure 5A**.

**Figure 5.**
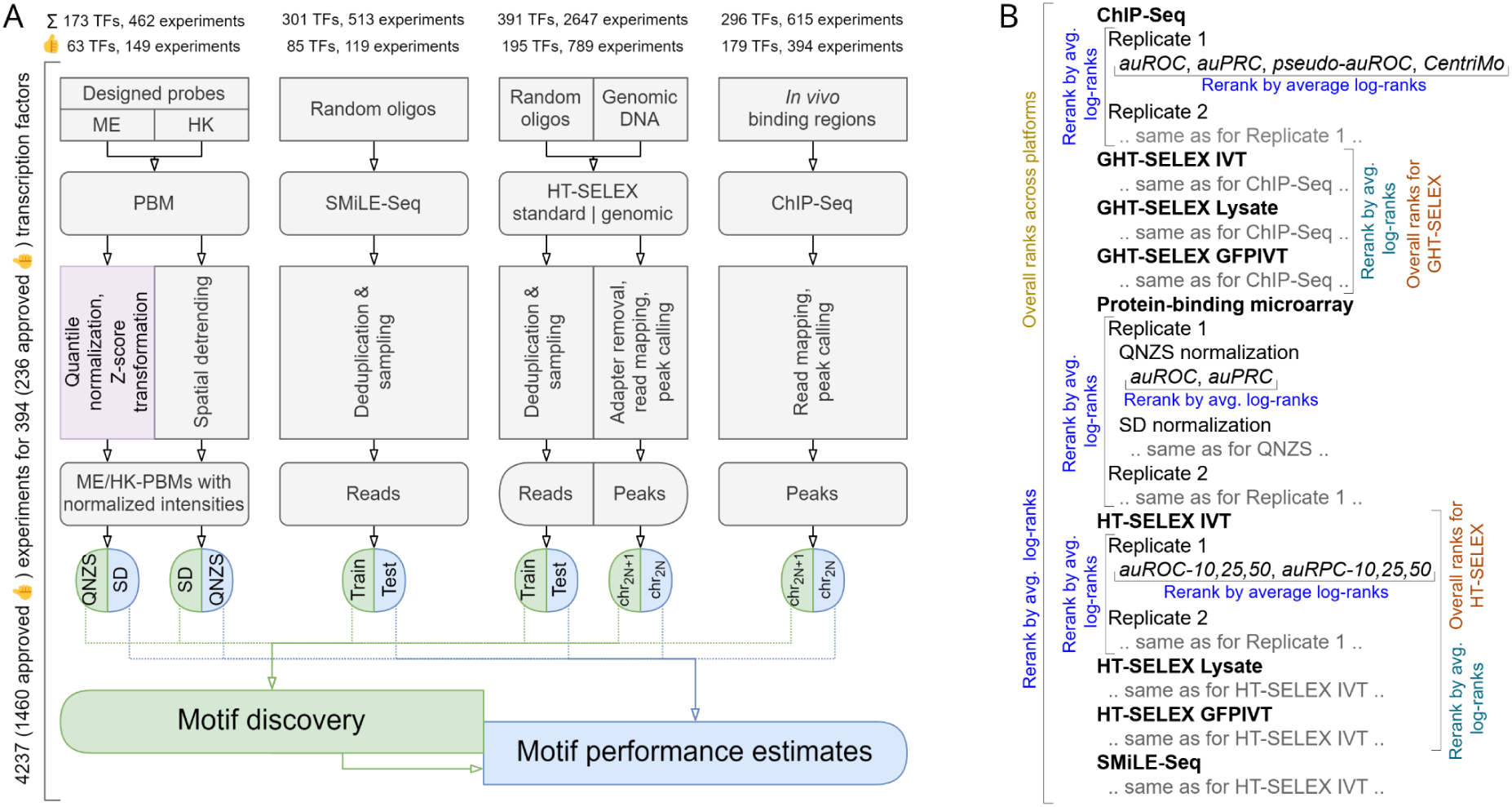
Schematics of the underlying workflows. A: Experimental data preprocessing and generation of train-test data slices for motif discovery and benchmarking. For (G)HT-SELEX, multiple SELEX cycles are counted as a single experiment. B: Scheme of the hierarchical rank aggregation.

#### HT-SELEX and SMiLE-Seq

No special preprocessing was performed for HT-SELEX or SMiLE-Seq data (FASTQ). However, in HT-SELEX and SMiLE-Seq, the binding sites may overlap the constant parts of the oligonucleotides that were physically present during the binding experiments, i.e., the binding sites may include parts of the primers and/or barcodes, which vary from experiment to experiment. Therefore, this information was saved in file names and metadata and then explicitly made available during motif discovery and benchmarking.

The sequence design of HT-SELEX data was the following: `5’ ACACTCTTTCCCTACACGACGCTCTTCCGATCT [BAR1] (40N) [BAR2] AGATCGGAAGAGCACACGTCTGAACTCCAGTCAC 3’` where BAR1 and BAR2 were experiment-specific barcodes, and 40N was a 40 bp random insert.

The sequence design of the Codebook SMiLE-Seq data was the following: `5’ CGTCGGCAGCGTCAGATGTGTATAAGAGACAG [BAR1] (40N)CTGTCTCTTATACACATCTCCGAGCCCA 3’` with a 40 bp random insert. The sequence design of previously published SMiLE-Seq data was the following: `5’ ACACTCTTTCCCTACACGACGCTCTTCCGATCT [BC-half1] (30N) [BC-half2] GATCGGAAGAGCTCGTATGCCGTCTTCTGCTTG 3’` with a 30 bp random insert.

#### Protein-binding microarrays

The Codebook data were obtained from two PBM designs, ME and HK^23^, which were preprocessed separately to account for systematic biases, for example, from the arrangement of probes on the microarray. We used two preprocessing strategies:

1. QNZS, quantile normalization followed by Z-scoring. Here log-transformed probe intensities underwent quantile normalization to make the signal distributions of each array identical. Next, the intensity of each probe underwent a Z-score transformation, with the mean and std. dev. assessed for each probe separately based on its intensities across all available PBMs.
2. SD, spatial detrending with window size 11×11 as tested in^19^. For motif discovery, we also performed quantile normalization after spatial detrending (SDQN) to have a uniform normalized scale for all datasets.

The software implementation of the procedures is available at https://github.com/autosome-ru/PBM_preprocessing.

#### ChIP-Seq and genomic HT-SELEX

The analysis of both ChIP-Seq and GHT-SELEX data was performed using the unified GTRD ChIP-Seq pipeline^52^. Both for ChIP-Seq and genomic HT-SELEX (GHT-SELEX, GHTS), the FASTQ read alignment was performed with *bowtie2* (2.2.3, default parameters and fixed `--seed 0`). For paired-end reads, we additionally specified `--no-mixed --no-discordant --maxins 1000`. Reported alignments were filtered by MAPQ score with *samtools* `-q 10`. For paired-end data, we additionally marked and removed PCR duplicates with *Picard MarkDuplicates*. Specifically, for the genomic HT-SELEX data, before read mapping, we performed adapter trimming with *cutadapt* (version 1.15, AGATCGGAAGAGC as the adapter sequence: `-a AGATCGGAAGAGC -A AGATCGGAAGAGC -o out.R1.fastq.gz -p out.R2.fastq.gz in.R1.fastq.gz in.R2.fastq.gz`).

To achieve a balanced sequencing depth between experiments and controls and reduce computational load, peak calling was performed against randomly sampled control data (10% of the total pooled set of control reads from the matching batch, sampling performed after the alignment step). For ChIP-Seq, the input DNA samples were used as the control. For genomic HT-SELEX, the zero-cycle unselected reads were used as control. Paired-end control data was prioritized for paired-end ChIP-Seq when available in the same batch.

For peak calling, four tools (*macs2* 2.1.2, *pics* https://github.com/Biosoft-ru/cpics, *gem* 2.5, *sissrs* 1.4) were executed with default settings, except for *macs2*. For the latter, for single-end reads, the expected fragment length `$frag_len` was estimated with a strand cross-correlation approach (*run_spp.R* from the ENCODE pipeline dated Aug 29, 2016, https://github.com/kundajelab/phantompeakqualtools). Next, *macs2* was executed with `--nomodel --extsize $frag_len` for single-end read alignments. For paired-end reads, we ran *macs2* in the paired-end mode (`-f BAMPE --nomodel`). Single-end peak callers (*pics*, *gem*, *sissrs*) were executed on paired-end data using alignments of the first reads in pair (*samtools* `-F 128 paired.bam`). The primary peak calls for each dataset were obtained with *macs2*. Next, technically reproducible *macs2* peaks were selected by ensuring a non-empty overlap with any of the peaks from other peak callers (*pics*, *sissrs*, *gem*). For GHT-SELEX, the peak calling was performed separately for reads originating from each cycle. The resulting files follow the *macs2* peak call format (narrowPeak) with an additional column listing supporting evidence from our peak callers. The resulting peaks were sorted by chromosome and coordinates.

### Train-test data splits and benchmarking protocols

In this study, we focused on a fair assessment of motif performance. Thus, for each experiment, we have generated separate non-overlapping training and test datasets. The only exception was PBM data, where we allowed a criss-cross training-test for different normalization strategies (SD and QNZS), i.e. for a particular PBM, QNZS data were applicable for testing SD-derived motifs and vice versa. The experiment-specific benchmarking train-test splits and protocols are described below. The implementation is available on GitHub (https://github.com/autosome-ru/motif_benchmarks). For all motif discovery tools, the resulting motif model was a matrix of positional nucleotide counts or a matrix of normalized nucleotide frequencies.

#### Benchmarking with PBM probe intensities

PBM data were used to assess motif performance using the binary classification of positives (specifically bound probes) and negatives (the rest of the probes): area under the receiver operating characteristic (auROC) and area under the precision-recall curve (auPRC) were computed with Jstacs^53^. For SD-preprocessed PBMs, the probes passing mean + 4 std. dev. intensity threshold were designated as positives as in the PBM-centric DREAM challenge of Weirauch et al.^19^, see Online Methods, “AUROC of probe intensity predictions” Section. For QNZS-normalized PBMs, probes with Z-scores above 4 were considered positives. In case these rules provided less than 50 positives, a minimum of 50 top-scoring probes was used instead. Motif scanning for this metric used the sum-occupancy scoring with PFMs. During scanning, the first 6 bps of the static linker sequence were concatenated to the unique sequence of each probe.

#### Benchmarking with ChIP-Seq and GHT-SELEX peaks

The train-test split for peaks data was performed using complete chromosome holdout: peaks at odd-numbered autosomes were designated for model training, and peaks at even-numbered autosomes for testing. Only experiments yielding 50 or more peaks in the test data were used for benchmarking, and only datasets with 50 or more peaks in the training data were used for motif discovery. Individual GHT-SELEX cycles were considered separately. Three different peak-based benchmarks were used.

1. The Orenstein-Shamir^16^ setup for binary classification of positives against neighboring negative regions yielding area under the receiver operating characteristic (auROC). We used the approach implemented in Ambrosini et al.^17^ with the following modifications: up to the top 1000 peaks were used to generate positives as 250bp-long [-124,+125] regions around the peak summits; for each positive peak, 250bp regions located 300bp upstream and downstream from the original peak summit were included in the negative set. Motif scanning for this metric used the sum-occupancy scoring with PFMs. The area under the precision-recall curve (auPRC) was estimated in addition to the auROC.
2. The asymptotic pseudo-auROC as in HOCOMOCO v11^54^. This method compared the top-scoring PWM motif hits in positives against the asymptotic estimate for random sequences of the same lengths and dinucleotide composition as in the positive sequence set. Positive regions’ lengths were standardized by taking ±150bp around the peak summits, and up to 1000 first peaks from the test data (reproducible peaks sorted by chromosome and coordinate, see above) were used. Motif scanning for this metric used best-hit log-odds scoring, and PFMs were transformed to log-odds PWMs in the following way^55^:

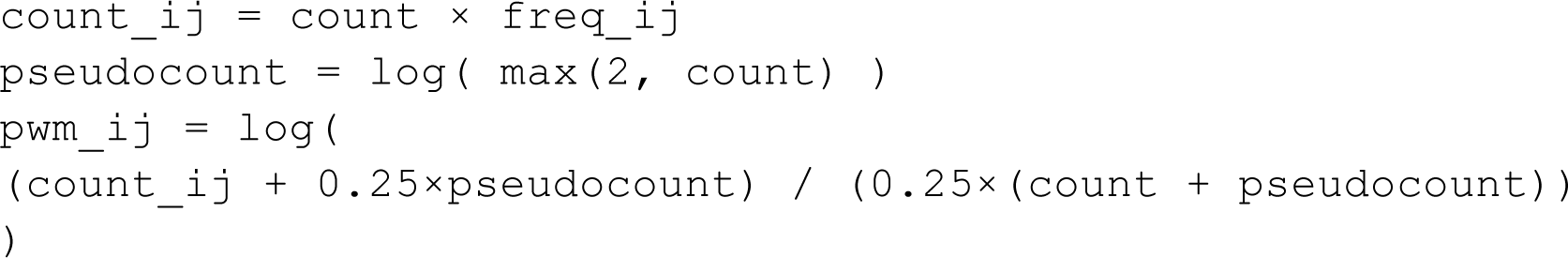

where `count_ij` is the *i,j*-th element of the matrix of non-normalized nucleotide counts. For ChIPMunk, `count` was set to the actual number of aligned words. For other methods yielding normalized PFMs, `count` was set to 100.
3. CentriMo^38^ motif centrality measure (-log-*E*-value) for motif hit locations against peak summits. For motif scanning, CentriMo performs the PFM-to-PWM transformation internally. In the case the run with default parameters technically failed to provide output (e.g. due to few sufficiently high-scoring sites), we reran CentriMo with `--score 1 --use-pvalues` to allow it considering low-scoring motif occurrences.

#### Benchmarking with HT-SELEX and SMiLE-Seq reads

For each cycle, reads were separated into the train and test datasets in a 2 to 1 ratio. At the benchmarking stage, the reads from different cycles (for HT-SELEX) were pooled together. We randomly sampled a maximum of 500,000 unique reads per dataset and used the Orenstein-Shamir benchmarking protocol (sum-occupancy scoring with PFMs) as in^17^ with 10%, 25%, or 50% of top-scoring reads to be designated as positives for each tested PWM. In addition to auROC (as in the original protocol), we also computed auPRC.

### Identifying best-performing motifs

For each motif and each test dataset, we computed several performance measures. To identify the best-performing motif, we performed hierarchical rank aggregation as suggested in the DREAM-ENCODE challenge (https://www.synapse.org/#!Synapse:syn6131484/). We ordered the motifs by achieved performance for all combinations of benchmarks and performance metrics and calculated the geometric mean of the ranks, followed by re-ranking (**Figure 5B**):

- first, across different metrics of a single benchmark (e.g. auROC and auPRC), then across variants of benchmarking settings (e.g. HT-SELEX benchmark with 10%, 25%, or 50% top hits taken as positives), and then across different benchmarks (e.g., for ChIP-Seq, Orenstein-Shamir classification performance and CentriMo motif centrality);
- next, across independent experiments of the same type (experimental replicates) and technical sequencing replicates, which were available for select ChIP-Seq datasets.

This provided the best motif for a TF for a particular type of experimental data. Next, aggregation across all data types was performed to identify the best motif for each TF in terms of the overall performance across experiment types.

The procedure was performed twice: once for the complete data (pre-curation) and once for the curation-approved datasets and respective motifs. The results of both variants are available online in Codebook MEX, and the curation-approved datasets were used for detailed analysis of motif performance in this study.

The software implementation of the ranking procedure is available on GitHub (https://github.com/autosome-ru/greco-bit-data-processing).

Note, that in the overall ranking (**Figure 1**), individual HT- and GHT-SELEX types (Lysate, IVT, GFPIVT) were considered as independent platforms, while in the subsequent analysis (**Figure 2**, **Figure 3**) they were considered together.

#### Harmonizing benchmarking measures

The range of performance measures depends not only on the benchmarking protocol but also on the TF and the particular experimental dataset. To make the benchmarking metrics comparable and to obtain a common scale in **Figure 2**, we applied a linear transformation to project the raw values of the benchmarking metrics (auROC, auPRC, etc) into the [0;1] range for each <TF, dataset, benchmark, metric> combination independently, where 0 corresponded to the lowest achieved value of the worst-performing motif, and 1 was the highest achieved value of the best-performing motif. At the cost of direct interpretability of metric score differences, this allowed quantitative analysis of metrics across TFs and experiment types.

### Dataset curation procedure

The availability of multiple types of experiments and replicates facilitated the comprehensive manual curation of datasets in terms of the consistency of DNA specificity patterns discovered from different types of experiments. In addition to the general consistency of motifs (e.g., visual similarity of logos) derived from different types of experiments, a major argument for dataset approval was the cross-experiment motif performance, i.e. motifs trained on one and high-scoring on the other type of experimental data were supporting approval of both the source and the benchmarking dataset. To simplify curation, we have annotated the motifs with the closest known patterns from HOCOMOCO v11^37^ using MACRO-APE^55^, this annotation is available in MEX from the curation stage.

During curation, each experiment was examined by at least one junior and two of three senior curators (A.J., I.V.K, and T.R.H.). The key features to analyze were the best-performing motifs, their datasets of origin, types of experimental data, and absolute achieved performance metrics. Cases with discordant approved/non-approved curation labels were individually rechecked, discussed, and resolved by two senior curators (A.J. and T.R.H.).

Of note, traditional quality controls, such as ENCODE-style metrics for ChIP-Seq data, were also taken into account during the curation and are available in the experiment metadata in MEX. However, there were cases when formally poor QC experiments yielded the proper motifs or ranked high proper motifs from other datasets, allowing approval of such datasets.

### Filtering artifact signals

The motif benchmarking and dataset curation could be complicated due to common artifacts related to particular types of experiments where highly similar motifs were observed in a large number of experiments for different proteins. Some of these artifacts, such as the ACGACG sequence observed in HT-SELEX, match constant flanking regions of the SELEX ligands and are likely derived from the enrichment of partially single-stranded ligands, whereas others are only seen in the lysate-based experiments and correspond to endogenous TFs (such as NFI and YY1) that are highly expressed in HEK293 cells.

To reduce the overall impact of these widespread artifact signals, we manually assembled a catalog of artifact motifs during the curation stage (see **Supplementary Table ST3**). Next, we filtered the whole motif collection by comparing the motifs against the catalog using MACRO-APE^55^ (with the motif P-value threshold 5·10^-4^ and `-d 10`). Motifs with Jaccard similarity ≥ 0.15 between high-scoring word sets^55^ were filtered out. We also filtered the motifs scoring below P-value 10^-4^ in constant flanks of HT-SELEX, genomic HT-SELEX, or SMiLE-Seq in a primer/barcode-dependent way. Of note, ETS-related motifs were not filtered for ETS-related “positive control” TFs (ELF3, FLI1, and GABPA).

### Motif discovery tools and data tool-specific data processing

The first round of motif discovery was focused on applying a diverse set of tools to the complete Codebook data, and its results were used for the dataset curation, as described above, to include only approved datasets in the downstream analysis of motif performance. The second round of motif discovery was to generate more motifs from the curation-approved datasets by employing popular motif discovery tools used in the first round with alternative settings and on more data types while including motifs yielded by the recently published ProBound software^39^. Of note, not all motif discovery tools were applicable in a ready-to-use fashion to all data types, and thus extra preprocessing was necessary to use the tools on data types, for which they were not designed in the first place.

#### ChIPMunk

ChIPMunk is a PWM-based greedy optimization algorithm suitable for processing thousands to tens of thousands of sequences with or without positional prior such as ChIP-Seq peak summit location.

##### Motif discovery from ChIP-Seq and GHT-SELEX peaks

###### Data preparation

ChIP-Seq and GHT-SELEX peak calls were sorted by peak height and top 500 and 1000 regions respectively were taken for subsequent analysis. 301-bp long regions centered on the peak summits were extracted for motif discovery in ChIPMunk “peak” mode by specifying 150 as the relative peak summit location.

###### Motif discovery

ChIPMunk launcher script was executed with the following parameters:

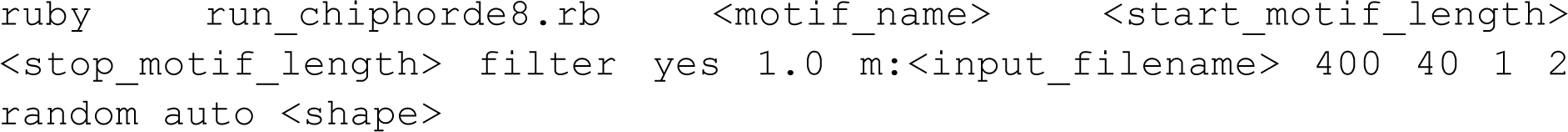

The motif discovery was performed three times using (1) 21 to 7 motif lengths range, no motif shape prior; (2) 15 to 7 lengths range, no motif shape prior; (3) 7 to 15 lengths range, “single-box” motif shape prior. For Zinc-finger TFs we expected longer motifs and used alternative settings: (1) 25 to 7 lengths range, no motif shape prior; (2) 7 to 21 lengths range “single-box” motif shape prior. Using the ‘filter’ strategy, ChIPMunk performs filtering of the initially found primary motif hits and could yield a secondary alternative motif.

##### Motif discovery from HT-SELEX, GHT-SELEX, and SMiLE-Seq reads

###### Data preparation

For (G)HT-SELEX, we pooled reads across all cycles of an experiment or for the terminal 3+ cycles only; the complete training data was used for SMiLE-Seq. To account for binding sites overlapping constant parts (technical segments) of the oligonucleotides, the reads were extended as 5’-NNX_1_<read>X_2_NN-3’, where X_1_ and X_2_ belonged to the technical segments and thus were the same across the sequences from a particular experiment. Singleton sequences found only once in the pooled dataset were excluded. Next, 5-mer enrichment against dinucleotide shuffled control was computed with the custom script (https://github.com/autosome-ru/HT-SELEX-kmer-filtering). For each dataset, we gathered 500, 1000, and 2500 top-enriched sequences for motif discovery with dinucleotide ChIPMunk^56^ and 10000 sequences for ChIPMunk; all available sequences were used in the case of fewer sequences than that number available. For dinucleotide ChIPMunk, the standard position weight matrices were constructed from the resulting multiple sequence alignments.

###### Motif discovery

The ChIPMunk was used via ruby launcher with the following parameters specifying 7 to 25 motif length range:

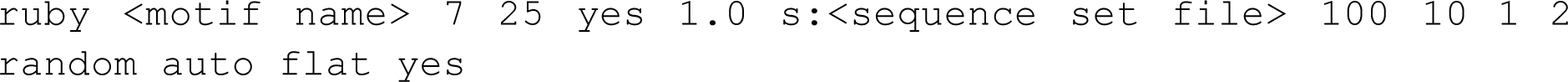

##### Motif discovery from PBM probes

###### Data preparation

For each microarray we used the sequences of the top 1000 probes ranked by normalized signal intensity, skipping the flagged probes^57^. The sequences were taken without the linker flank.

###### Program execution

The dinucleotide version of ChIP-Munk was used with the following parameters:

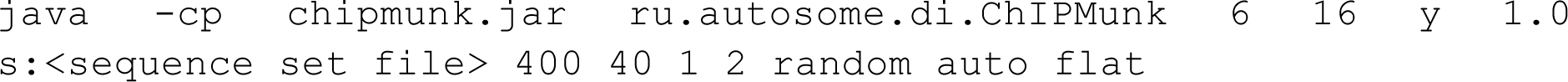

The single-nucleotide position count matrices were constructed from the multiple sequence alignments.

#### Dimont

Dimont is a motif discovery algorithm that allows for modeling binding motifs using Markov models in general and PWMs in particular. Dimont has been designed for using all available sequences from binding experiments (e.g., all ChIP-Seq peaks), where each sequence is associated with a measure of confidence that this specific sequence is bound (e.g. peak statistics from ChIP-Seq experiments), which are converted to soft class labels (bound vs unbound) with assay-specific formulas. The objective function of Dimont (maximum supervised posterior) optimizes the concordance of these soft labels and motif-based scores using gradient-based numerical optimization, i.e., Dimont tries to find the motifs that explain the soft labels best. Dimont-HTS is a variant of the Dimont algorithm with an HTS-specific weighting schema for the soft labels and an adapted initialization strategy.

Motif discovery using Dimont requires an input set of sequences, which are complemented by a sequence-specific “signal” annotation, which indicates the confidence that a specific sequence is bound by the TF of interest. Signal values are converted to soft labels internally using a rank-based method^33^, where a value of 1 indicates perfect confidence that a sequence is bound and 0 indicates perfect confidence that a sequence is not bound by the TF. Dimont then aims at finding the motif that explains the soft labels best, i.e., that yields high scores for sequences with soft labels close to 1 and low(er) scores for sequences with soft labels close to 0. This is achieved by maximizing the supervised posterior of the soft-labeled input data^33^. In the following, we describe for the specific data types how sequences were extracted, how “signal” values were defined, and how these were used for motif discovery using Dimont.

##### Data preparation for ChIP-Seq and GHT-SELEX peaks

All ChIP-Seq and GHT-SELEX peaks in the training set were considered and 1000-bp long regions around the peak centers were extracted together with the corresponding peak statistics (column 7 of the peak list) and stored in FastA format. The peak statistic was used as a “signal” annotation in the FastA headers of the extracted sequences and is subsequently used for determining weights in the Dimont learning procedure.

##### Data preparation for PBM probes

For each probe, the unique probe sequence and the first 6 bp of the static linker sequence were concatenated and extracted together with the mean signal intensity value of the corresponding probe and stored in FastA format. The mean signal intensity was used as a “signal” annotation in the FastA headers of the concatenated sequences.

##### Data preparation for HT-SELEX reads

First, reads from all HT-SELEX cycles were extracted and stored in FastA format using the HT-SELEX cycle as a “signal” annotation. Then reads across all cycles of an HT-SELEX experiment for a specific TF were sub-sampled to at most 400,000 reads, while sampling reads from the different cycles such that the distribution across cycles was as even as possible.

##### Data preparation for SMiLE-Seq reads

Reads from each SMiLE-Seq experiment were extracted and stored in FastA format. For a specific SMiLE-Seq experiment, all reads were assigned a “signal” annotation of 1. These were complemented with a sub-sample of one-fifth of the original reads from all other SMiLE-Seq experiments from the same batch (barcode) but for other target TFs, which were assigned a “signal” annotation of 0.

##### Motif discovery from ChIP-seq and GHT-SELEX peaks, PBM probes, and SMiLE-Seq reads

For sequences generated from ChIP-seq and GHT-SELEX peaks, PBM probes, and SMiLE-Seq reads, Dimont was executed with default parameters with a few minor exceptions. For ChIP-seq and GHT-SELEX, the initial motif width was set to 20 (imw=20). For PBM probes and SMiLE-Seq, the initial motif width was set to 10, and masking of previous motif occurrences was switched off (`imw=10 d=false`), while the weighting factor was set to 0.05 (`w=0.05`) for PBM probes and 0.5 (`w=0.5`) for SMiLE-Seq. The first two motifs reported by Dimont were used for further analyses.

##### Motif discovery from HT-SELEX reads

For HT-SELEX, two alternative strategies were used. In the first approach, the cycles stored as “signal” values were converted to soft labels based on an enrichment factor E as E^cycle-max(cycle)^. Aside from the definition of soft labels, Dimont was started with default parameters. The second approach (Dimont-HTS) was more specifically tailored to HT-SELEX data. Here, the motif initialization step of Dimont was based on 10-mers identified by a re-implementation of the Z-score proposed by Ge et al.^58^ and filtered for redundancy using a minimum Huddinge distance^31^ of 2. The determination of soft labels was based on a cycle-specific and a sequence-specific weight interpolating linearly between adjacent cycle-specific weights. The cycle-specific weight was determined from the relative number of unspecific sequences in each HT-SELEX cycle using a re-implementation of the method proposed by Jolma et al.^15^. Within each cycle, sequence-specific weights were determined based on the ranks of 8-mer occurrences among all sequences of a cycle. Sequence-specific weights were defined as the maximum relative rank of an 8-mer occurring in a sequence divided by the maximum rank across all sequences. Besides the adapted initialization strategy and determination of soft labels, the initial motif width was set to 20 (`imw=20`). Again, the first two motifs reported by Dimont were used for further analyses. The respective code is available at GitHub (https://github.com/Jstacs/Jstacs/tree/master/projects/dimont/hts).

#### ExplaiNN

ExplaiNN is a fully interpretable and transparent sequence-based deep learning model for genomic tasks that combines the powerful pattern recognition capabilities of convolutional neural networks with the simplicity of linear models.

##### Data preparation

To construct the training and validation datasets, each experiment was processed separately. Additionally, for the PBM data, we avoided mixing data from different normalization methods. As a data augmentation strategy, we doubled the size of each training and validation set by including the reverse complement of each sequence.

###### ChIP-Seq data

The peaks were resized to 201 bp by extending each peak summit by 100 bp in both directions. Then, they were randomly split into training (80%) and validation (20%) sets using the `train_test_split` function from scikit-learn (version 0.24.2, random splits were always performed in this manner)^59^. To avoid the need for negative samples during training, we retained the peak heights associated with each peak, thereby converting the training process into a regression task.

###### HT-SELEX and GHT-SELEX data

We treated cycles as independent classes, following the approach used by Asif and Orenstein^60^, thereby avoiding the need for negative samples during training. The reads were then randomly split into training (80%) and validation (20%) sets while maintaining the proportions between reads from each cycle.

###### PBM data

The probes, including both the de Bruijn and linker sequences, were randomly split into training (80%) and validation (20%) sets. Since the training task involved regression (see below), negative samples were not required.

###### SMiLE-Seq data

A set of negative samples was obtained by dinucleotide shuffling using BiasAway^61^ (version 3.3.0). Then, the original reads (positives) and the negative samples were combined and randomly split into training (80%) and validation (20%) sets, ensuring an equal proportion between positives and negatives.

##### Model training

All models featured the same architecture: 100 units and a filter size of 26. They were trained using the Adam optimizer^62^ for a maximum of 100 epochs. An early stopping criterion was set to halt training if the validation loss did not improve after 10 epochs. We applied one-hot encoding to the input sequences, converting nucleotides into 4-element vectors (*i.e.* A, C, G, and T). The learning rate was set to 0.003, and we used a batch size of 100. During training, we employed three different loss functions, tailored to each data type.

###### ChIP-Seq data

ExplaiNN was configured to model the peak heights using the negative log-likelihood loss with a Poisson distribution of the target (*i.e.* `PoissonNLLLoss` class from PyTorch^63^).

###### HT-SELEX, GHT-SELEX, and SMiLE-Seq data

The modeling tasks for these data involved either multi-label classification (*i.e.* for SELEX) or binary classification (*i.e.* for SMiLE-Seq). As a result, we chose `BCEWithLogitsLoss` as the loss function (*i.e.* binary cross-entropy with sigmoid).

###### PBM data

ExplaiNN was applied to model normalized intensity signals, making the mean squared error (MSELoss) the appropriate choice for the loss function.

##### Motif discovery

Following the specifications from the ExplaiNN manuscript, for each model, we constructed a position frequency matrix (PFM) for each filter by aligning all 26-mers (*i.e.* 26 bp-long DNA sequences) activating that filter’s unit by ≥ 50% of its maximum activation value in correctly predicted sequences. Then, we transformed the resulting PFMs into position weight matrices (PWMs), setting the background uniform nucleotide frequency to 0.25, and clustered them based on their Tomtom similarity^64^ using scripts from (https://github.com/vierstralab/motif-clustering). Finally, for each experiment (*i.e.* for each model), we returned the top 5 non-redundant PWMs (*i.e.* belonging to different clusters) based on their performance on the corresponding validation set.

#### GkmSVM with GkmExplain

##### Data preparation

ChIP-Seq data were sorted based on the q-value and the top 5000 peaks were taken. The top 5000 peaks were split into training and testing based on chromosomes (chr1 and chr3 used for testing). The peaks were extended by 100 bps on each side of the summit. For the negative set, for training, we used the `fasta-dinucleotide-shuffle-py3.in` from MEME suite^28^ to generate dinucleotide shuffled peaks from our positive set data.

##### Model training

To train the gkmSVM model^51^ we used the gkmtrain function from the LS-GKM package using the default parameters. LS-GKM is a version of gkmSVM^36^, an SVM-based algorithm that utilizes gapped k-mers as features. LS-GKM is specifically optimized for processing and training on a large number of sequences efficiently. In the default parameters, the word length (`-l`) is 11, the gap (`-d`) is 3 and the kernel used is the center-weighted (wgkm) kernel.

##### Motif discovery

To generate motifs we first generated importance scores and hypothetical importance scores using GkmExplain^36^. GkmExplain is a feature attribution technique applied to trained gkmSVM models that use a modified version of the integrated gradients method to determine the importance of individual nucleotides for the output label. GkmExplain has been shown to outperform^36^ other feature attribution methods such as deltaSVM^65^ and in-silico mutagenesis (ISM)^66^. Importance scores were generated from the test sequences and the train gkmSVM model using the command `gkmexplain` from LS-GKM package. The hypothetical importance scores were generated using the same command but with the parameter `-m 1`. To generate motifs from these importance scores we ran TF-MoDISco (https://github.com/kundajelab/tfmodisco) with the following parameters (`target_seqlet_fdr=0.2, sliding_window_size=21, flank_size=10, min_passing_windows_frac=0.0005`). TF-MoDISco uses importance scores derived from feature attribution methods to identify regions of high importance across sequences and clusters these recurring regions to generate motifs. Therefore, gkmExplain coupled with TF-MoDISco can be used to generate motifs from k-mer-based SVM models trained on our assays.

#### HOMER

Homer is a motif discovery algorithm that uses word enumeration followed by the hypergeometric or binomial test to detect oligo enrichment in the input sequence^29^. HOMER then transforms the sets of detected oligos into PWMs via an iterative refinement and optimization process.

##### Data preparation

ChIP-Seq, HT-SELEX, GHT-SELEX, and SMiLE-Seq data were processed in the same way as for ChIPMunk (see above).

##### Motif discovery

We called the `findmotifs.pl` function with default parameters to find motifs using HOMER for all experimental assays. For the negative set required by HOMER we generated dinucleotide shuffled using the `fasta-dinucleotide-shuffle-py3.in` script from MEME suite^28^. We also ran `findmotifs.pl` to find longer motifs up to 30bps by changing the `-len` parameter. The top 5 motifs outputted by HOMER for each set of parameters were used for analysis.

#### MEME

The Multiple EM for Motif Elicitation (MEME) employs the expectation maximization (EM) technique to derive PWMs. The algorithm begins by detecting an initial seed motif, which is then iteratively optimized through EM steps, which continue until the PWM values stabilize or a predefined iteration limit is reached. MEME primarily operates using the Zero or One Occurrence Per Sequence model to discover ungapped motifs of fixed lengths.

##### Data preparation

Data for Chip-Seq, HT-SELEX, GHT-SELEX, and SMiLE-Seq was processed in the same way as for ChIPMunk (see above).

##### Motif discovery

We ran MEME-ChIP^67^ on ChIP-Seq data and MEME^28^ on data from other assays. Both MEME-ChIP and MEME were first run using default parameters. We also ran both MEME-ChIP and MEME using `--maxw 30` and `--minw 3` to account for the longer motifs of C2H2 Zinc-Finger TFs. The top 3 motifs outputted by MEME for each set of parameters were used in the downstream analysis.

#### RCade

The Recognition Code-Assisted Discovery of Regulatory Elements (RCADE) algorithm is specifically built to uncover the binding preferences of the largest family of human transcription factors, the C2H2 zinc-fingers proteins. By utilizing predictions from the DNA recognition code specific to Zinc Fingers^68^, RCADE effectively infers the predicted binding motifs that are enriched in peaks compared to shuffled sequences.

##### Data preparation

Data for Chip-Seq, HT-SELEX, GHT-SELEX, and SMiLE-Seq was processed in the same way as for ChIPMunk (see above).

##### Motif discovery

We used RCADE2^35^ (https://github.com/csglab/RCADE2) using default parameters to identify motifs for C2H2-Zinc Finger TFs across all the assays. The amino acid sequences of the entire TF used as a parameter by RCADE2 were downloaded from UniProt. The top motif outputted by RCADE2 was used in the subsequent analysis.

#### STREME

STREME operates using a generalized suffix tree, a data structure similar to those used by tools like HOMER. STREME utilizes suffix trees to efficiently store input sequences and count matches between candidate PWMs (instead of oligos like Homer) and these sequences. After identifying potential motifs, STREME evaluates their enrichment in the input sequences using a one-sided Fisher’s exact test against control sequences. Like MEME, STREME operates under the assumption of a Zero or One Occurrence Per Sequence (ZOOPS) model.

##### Data preparation

Data for Chip-Seq, HT-SELEX, GHT-SELEX, and SMiLE-Seq was processed in the same way as for ChipMunk (see above).

##### Motif discovery

We ran STREME^32^ on data from all assays with two different sets of parameters. For the first run, STREME was run with default parameters. STREME was also run using `--maxw 30` and `--minw 3` to account for the large motif size of C2H2 Zinc-Finger TFs. The top 3 motifs outputted by STREME for each set of parameters were used for analysis

#### ProBound

##### Data preparation

For motif discovery using ProBound, k-mer count tables for each experiment were generated using all sequencing reads. The k-mer length was set to the entire probe length for SMiLE-Seq and HT-SELEX experiments. For GHT-SELEX, the probes were centered before extracting 40bp of sequence (20bp up and downstream of the center). Reads shorter than 40bp were discarded. In the case of multi-round SELEX experiments, columns indicating the round of enrichment were added to each count table. Since ProBound requires a sample of probe counts for an unselected input library, models could only be fit to SMiLE-Seq, GHT-SELEX, and HT-SELEX experiments. For SMiLE-Seq, input data was readily available for each experiment. For GHT-SELEX, input libraries were not matched to respective samples. Therefore, all input libraries were pooled and 10,000 unique reads were sampled at random to build a global input count table. For HT-SELEX, a deeply sequenced input library was unavailable. To approximate the input library, the reads from ‘failed’ experiments (those that showed no reliable probe enrichment after incubation with TFs) were combined, and 10,000 unique reads were sampled to create an approximate input count table. Note, that this approach is not recommended in the original publication and may bias motif inference, e.g., the approximate input is nonetheless subject to non-specific binding preferences.

##### Motif discovery

ProBound was used with the following, default optimizer settings: L2 regularizer weight of 0.000001 (`lambdaL2` parameter), Dirichlet regularizer weight of 20 (`pseudocount` parameter), smallest improvement in likelihood required for a model variation to be accepted of 0.0002 (`likelihoodThreshold` parameter). All values were taken from the ProBound documentation of single-experiment transcription factor binding models. Other optimizer settings were left at default values and no custom optimization was performed. Each experiment was analyzed with a single position-specific affinity matrix, which represents the change in binding affinity (K_d_) for all point mutations with respect to the optimal reference sequence^69^ binding mode with an initial size of 12 base pairs. For each experiment, a pair of models was developed - one incorporating the non-specific binding mode and the other one excluding it. In order to comply with the benchmarking pipeline, the energy logos produced by ProBound were first converted to position-specific affinity matrices (PSAMs) and then scaled to represent PFMs.

#### Autoseed

Autoseed generates motifs with two sequence sets, e.g., for HT-SELEX it uses a “signal” cycle (e.g. cycle3 of an experiment) and a “background” cycle (e.g. cycle 2), setting an IUPAC base sequence (e.g. ACCGGAAGRN) as a seed and then obtaining a motif based on this sequence and all sequences that are within a parameter specified edit distance from the seed (1, 2 or 3 edits). As in the previous work^31^, Autoseed was used to find Huddinge Distance-based local maxima for gapped 8-mers for combinations of a background and a signal cycle and to generate logos for these and heatmaps that display all possible spacing variants. Final motifs were generated manually by examining the Autoseed outputs to select optimal input parameters for motif generation.

### Random Forest of PWMs

#### Generating positive and negative data sets

We specially adapted the MEX data to allow for unbiased training and testing of advanced models suitable for genomic TFBS prediction. To enable testing the transferability of predictions between data types, we selected 142 TFs with at least one approved ChIP-Seq and at least one approved GHT-SELEX experiment. Separately for ChIP-Seq and GHT-SELEX, for each TF, the available peak sets were merged, and peak summit locations of the overlapping peaks were averaged. The choice of train-test chromosome hold-out was the same as in the primary MEX benchmarking.

The ‘positive’ sets (the bound regions) were created by extracting 301bp-long regions centered at the resulting peak summits. Three alternative negative sets were generated:

1. *Random* genomic regions (1:100 positive-to-negative class balance). Random regions were sampled from the genome matching the GC content distribution of the positive dataset.
2. *Alien* peaks of other TFs (1:100 balance) were also sampled and extracted in the same manner to match the GC content distribution of the positive set. In the case of TFs with larger positive sets, all available peaks were taken without GC matching if the 1:100 ratio was technically unachievable.
3. *Shades* of true positive peaks, the neighboring upstream and downstream regions (upto 1:2 balance). For each positive peak, the summit of a fake upstream peak was uniformly selected from [-750bp, -450bp] interval relative to the true positive summit, and the summit of a fake downstream peak was uniformly selected from [450bp, 750bp] interval. In the end, the achieved balance was often closer to 1:1 as the regions to sample the shades were overlapping blacklisted regions (see below) or peaks of the same transcription factor.

For all types of negatives, we explicitly excluded positive regions (whole peaks), ENCODE blacklist regions^70^, and any genomic regions with N nucleotides. The resulting set of TFs included 140 TFs, as the initially included CAMTA2 and FLYWCH1 were discarded due to having fewer *shades* than *positives*.

#### The ArChIPelago model

ArChIPelago, the arrangement of multiple position weight matrices with ChIP-Seq and machine learning for prediction of transcription factor binding sites, is a random forest model built on top of multiple PWMs. To construct Archipelago, separately for GHT-SELEX and ChIP-Seq data for each TF, we used the top 20 MEX PWMs best-performing at each replicate of ChIP-Seq and GHT-SELEX. ChIP-Seq-derived PWMs were not considered when training the model on GHT-SELEX and vice versa to prevent information leakage when evaluating model transferability.

The PWM predictions, i.e. the features for building the Random Forest, were obtained with SPRY-SARUS^71^: the log-odds PWMs best hits in each sequence were identified using `--skipn --show-non-matching --output-scoring-mode score besthit`. The resulting feature matrix with class labels (1,0) was scale-transformed with `sklearn.preprocessing.StandardScaler` from scikit-learn 1.3.2, and used to train a Random forest classifier model with the following hyper-parameters: `[’max_depth’: 6, ‘max_samples’: 0.8, ‘n_estimators’: 100]`. The *random* negative set was used for model training. To estimate the Archipelago performance, we computed auROC and auPRC with PRROC R package^72^ with three alternative negative datasets to reliably measure the model prediction quality.

## Data and Code Availability

The interactive Codebook/GRECO-BIT Motif Explorer website is available online at https://mex.autosome.org. The complete set of MEX motifs and the benchmarking-ready Codebook data are available at ZENODO^40–42^. The benchmarking protocols are available on GitHub (https://github.com/autosome-ru/motif_benchmarks). The implementation of the data processing pipeline is available on GitHub (https://github.com/autosome-ru/greco-bit-data-processing).

The software tools used in the study are listed in **Supplementary Table ST4**. The code supporting Random Forest model training and validation is available on GitHub (https://github.com/autosome-ru/MEX-ArChIPelago), and the respective data are available on ZENODO^73^.

## Acknowledgments

We wholeheartedly thank the IT Group of the Institute of Computer Science at Halle University for computational resources and personally Maximilian Biermann for valuable technical support. We thank Gherman Novakovsky for providing useful feedback, Debashish Ray for assistance with database depositions, and Irina Eliseeva and Valery Vyaltsev for their help with the preparation of the graphical abstract. We thank members of the GRECO consortium and personally Martin Kuiper for the encouragement and support of this project at its early stage in the form of dedicated workshops of the GREEKC COST action.

