## Supplementary figures and images for "Cross-platform DNA motif discovery and benchmarking to explore binding specificities of poorly studied human transcription factors"

### Supplementary Figure SF1

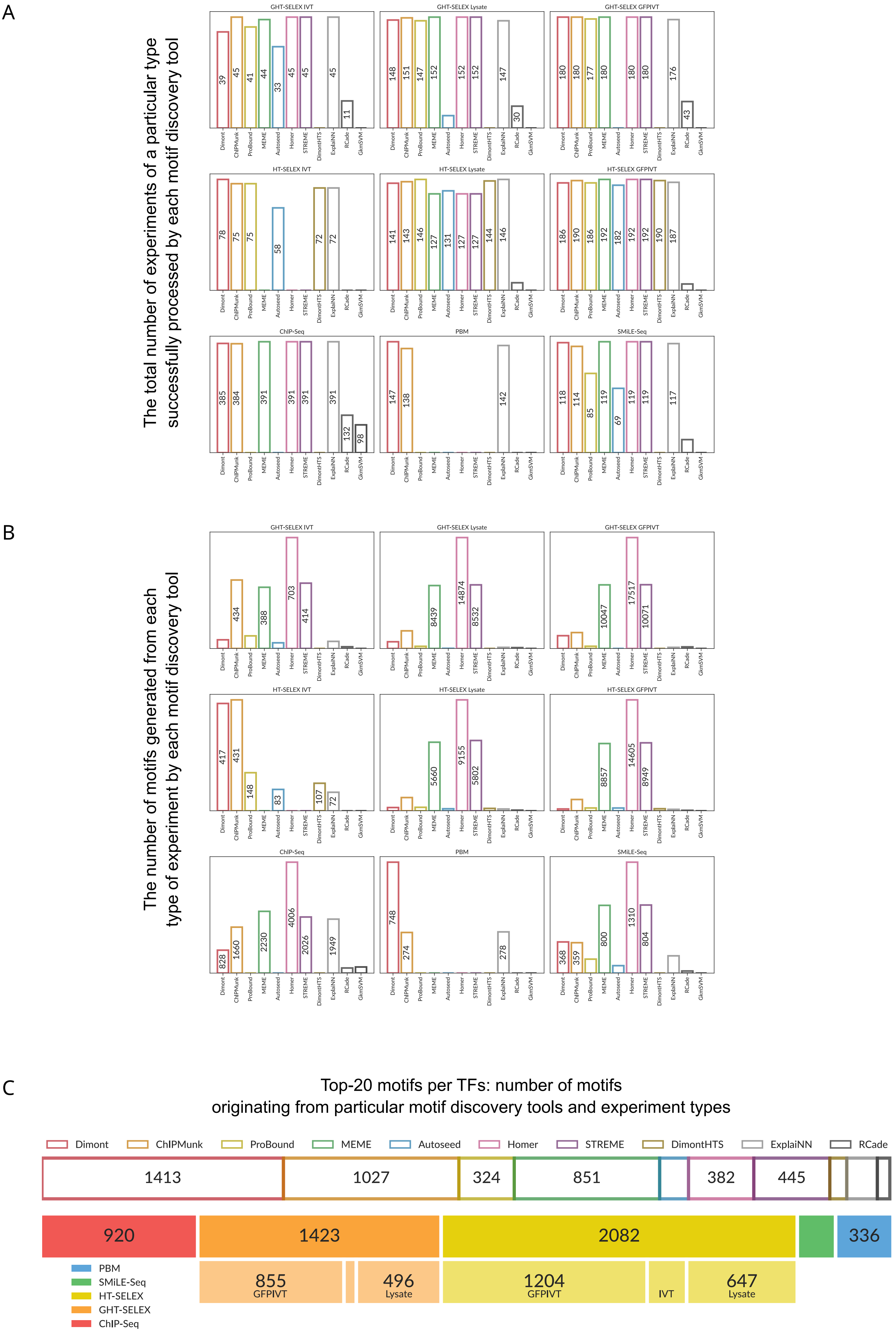


Supplementary Figure SF1

### Supplementary Figure SF2

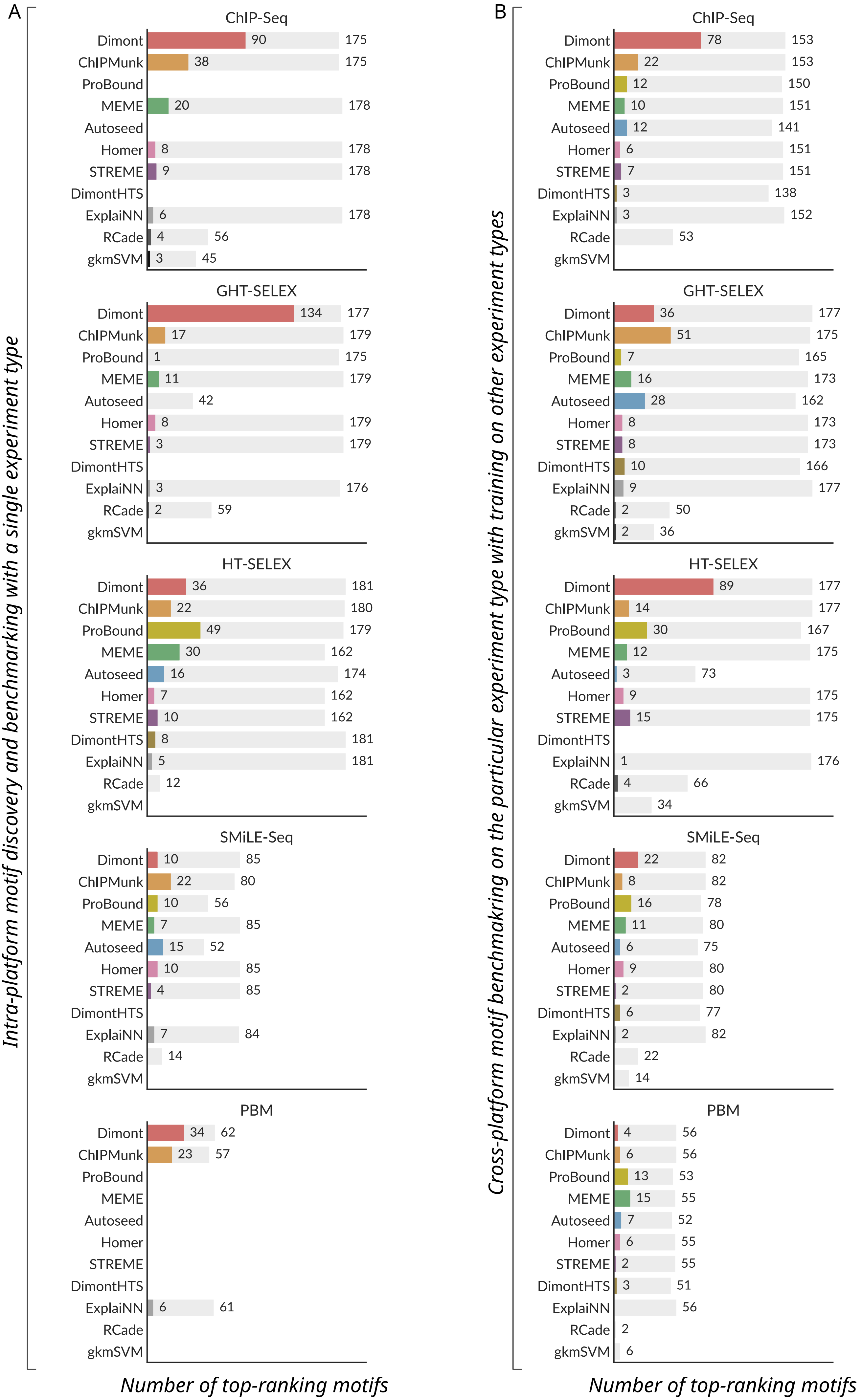


*grey bars denote the total numbers of eligible TFs in each category

Supplementary Figure SF2

### Supplementary Figure SF3

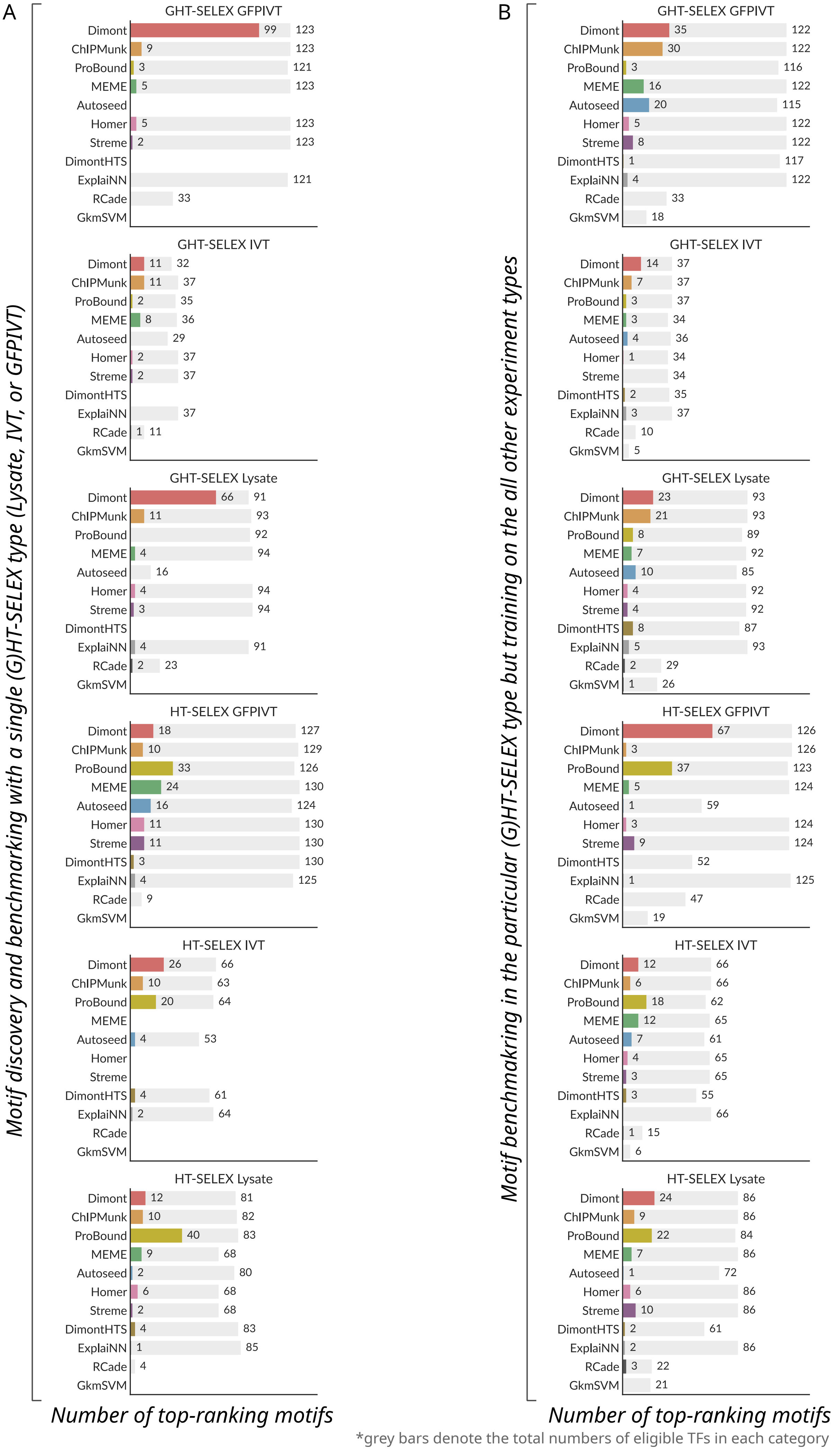


*grey bars denote the total numbers of eligible TFs in each category

Supplementary Figure SF3

### Supplementary Figure SF4

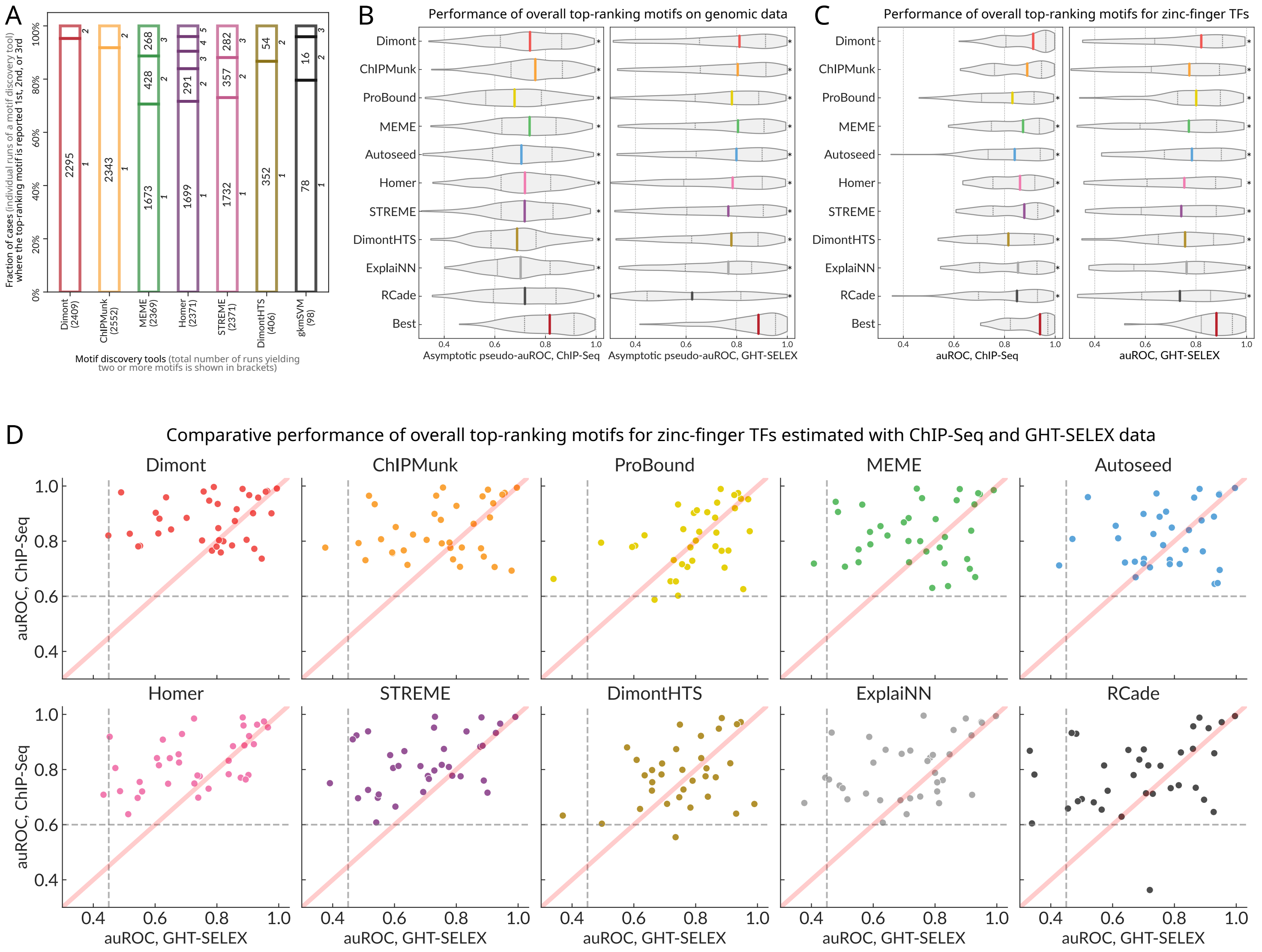


Supplementary Figure SF4

### Supplementary Figure SF5

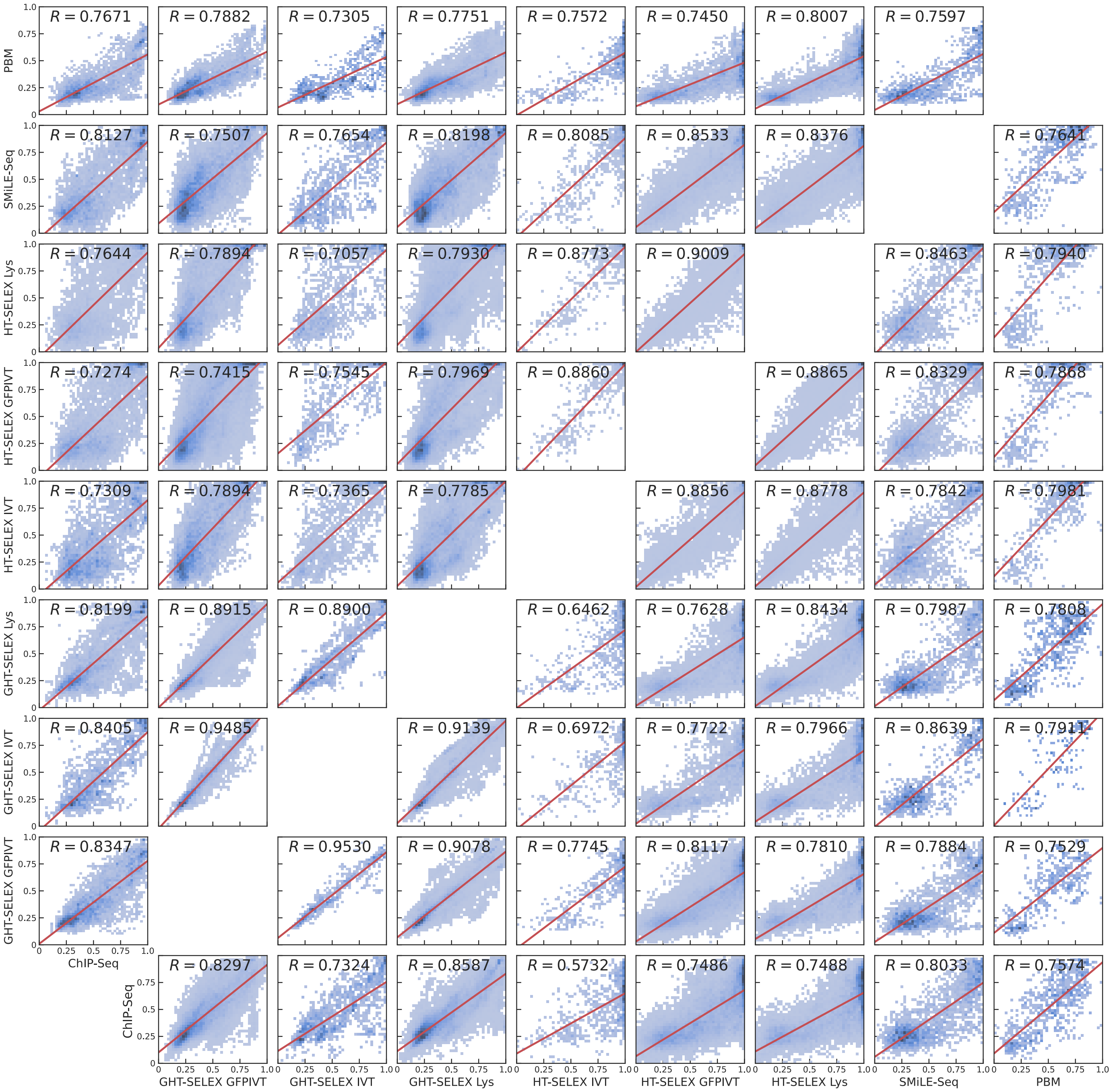


Supplementary Figure SF5

### Supplementary Figure SF6

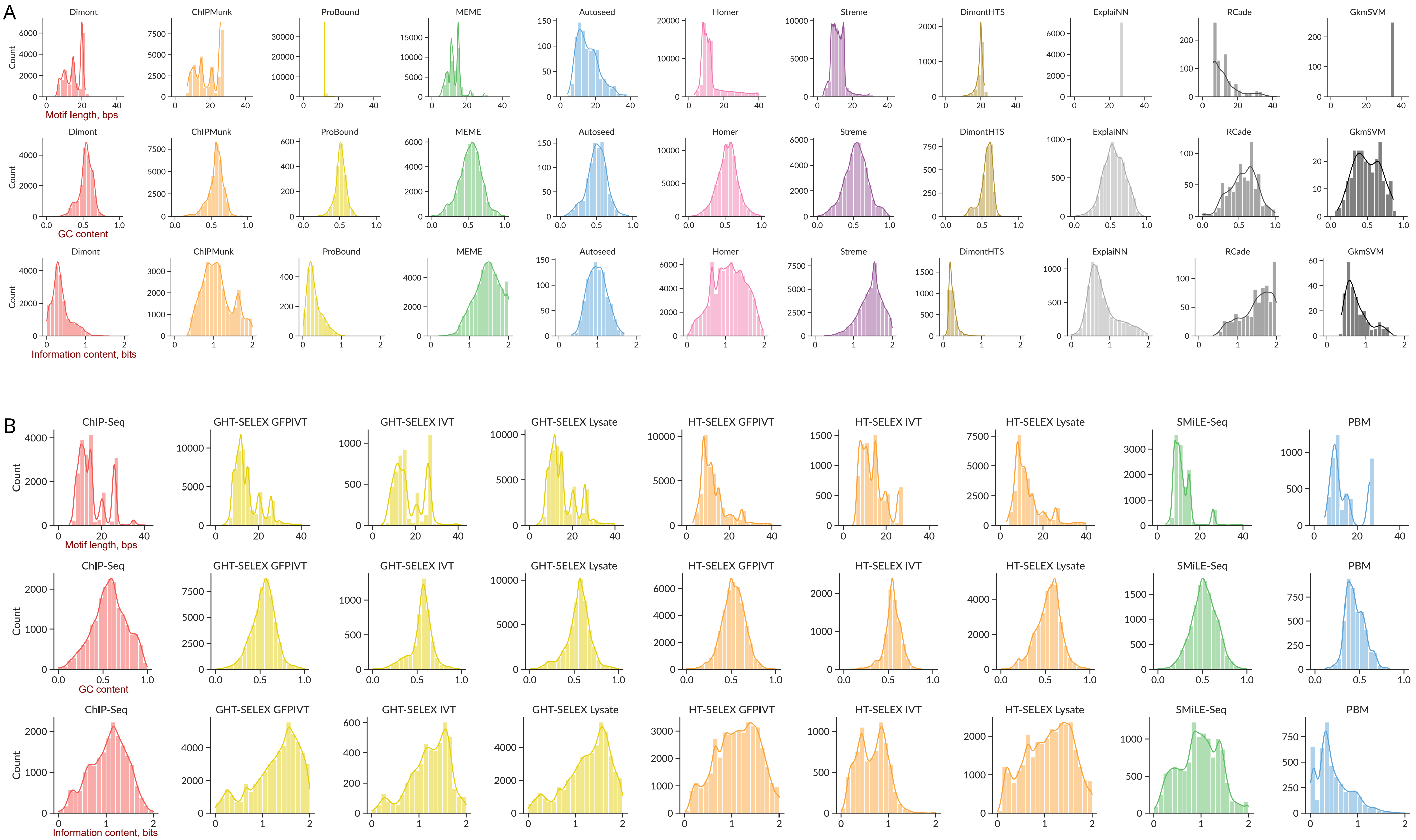


Supplementary Figure SF6

### Supplementary Figure SF7

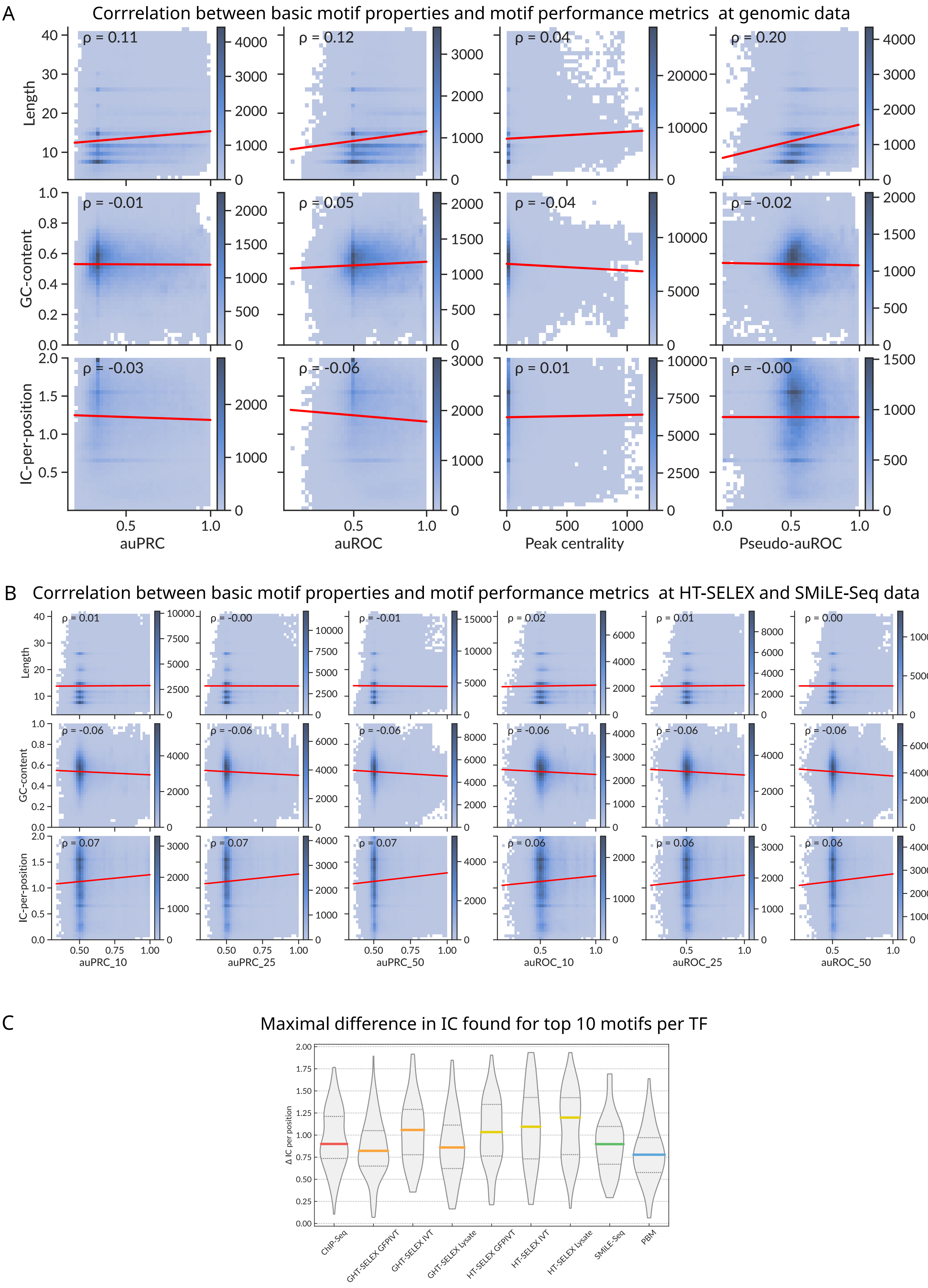


Supplementary Figure SF7

### Supplementary Figure SF8

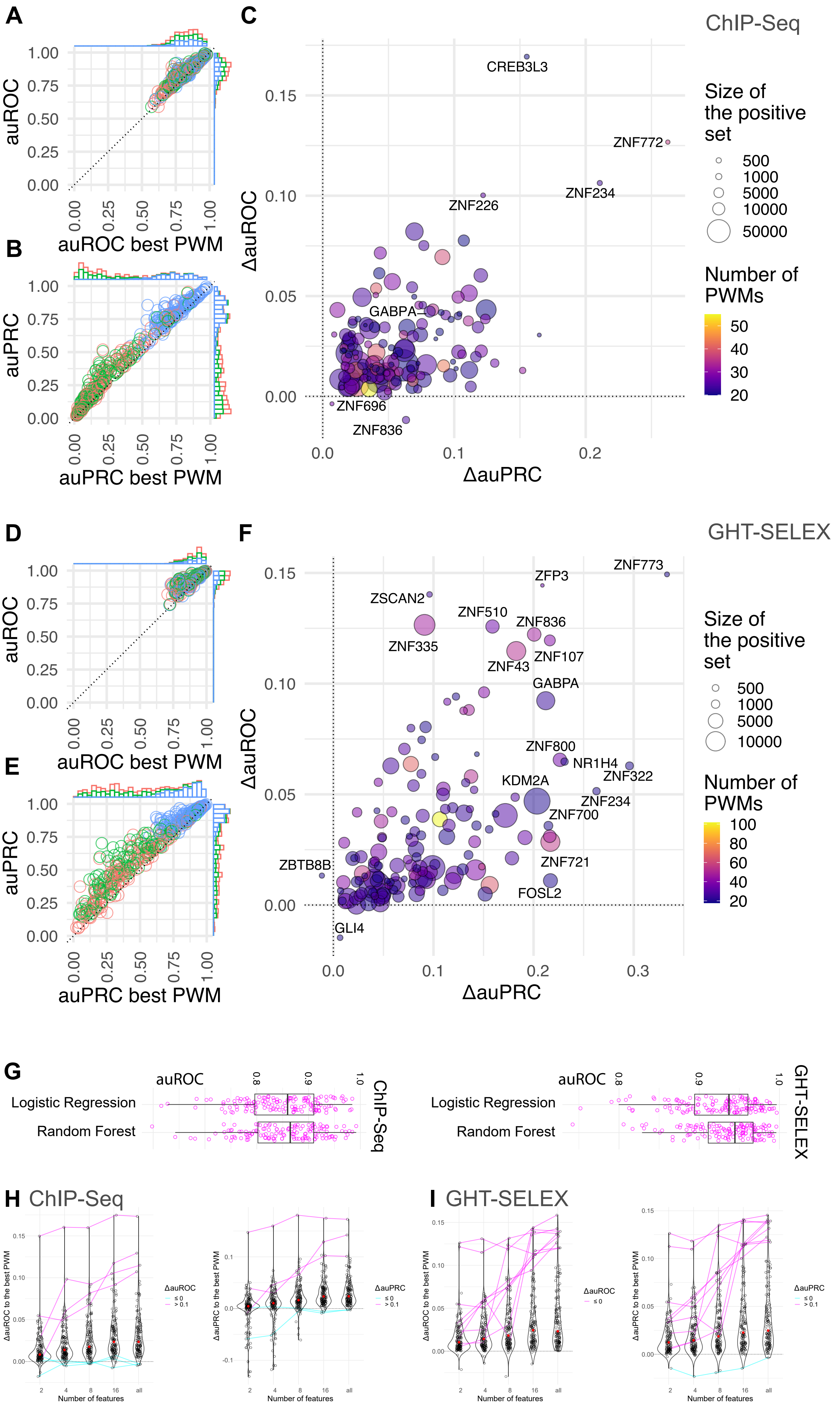


Supplementary Figure SF8

### Supplementary Figure SF9

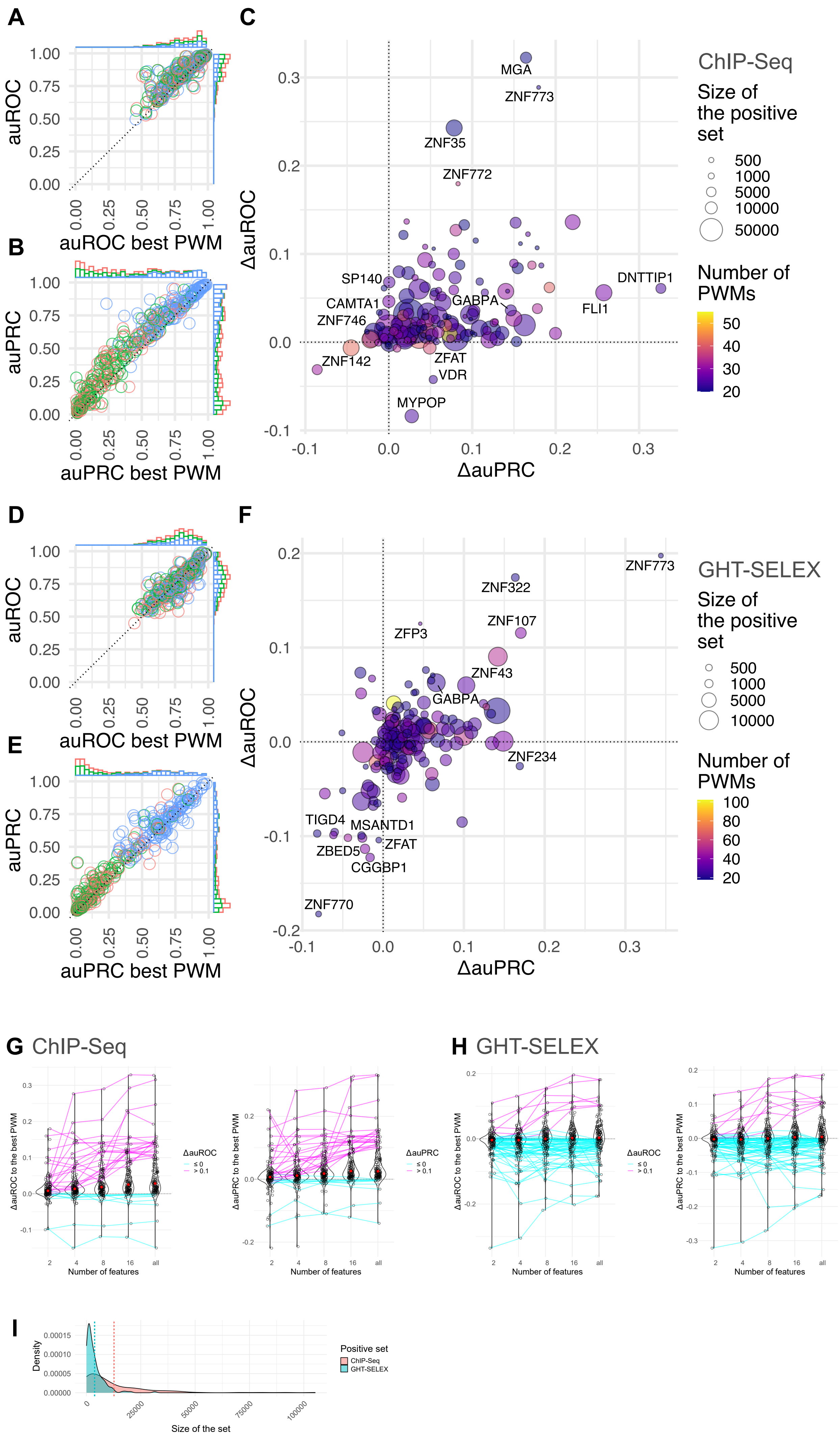


Supplementary Figure SF9

### Supplementary Figure SF11

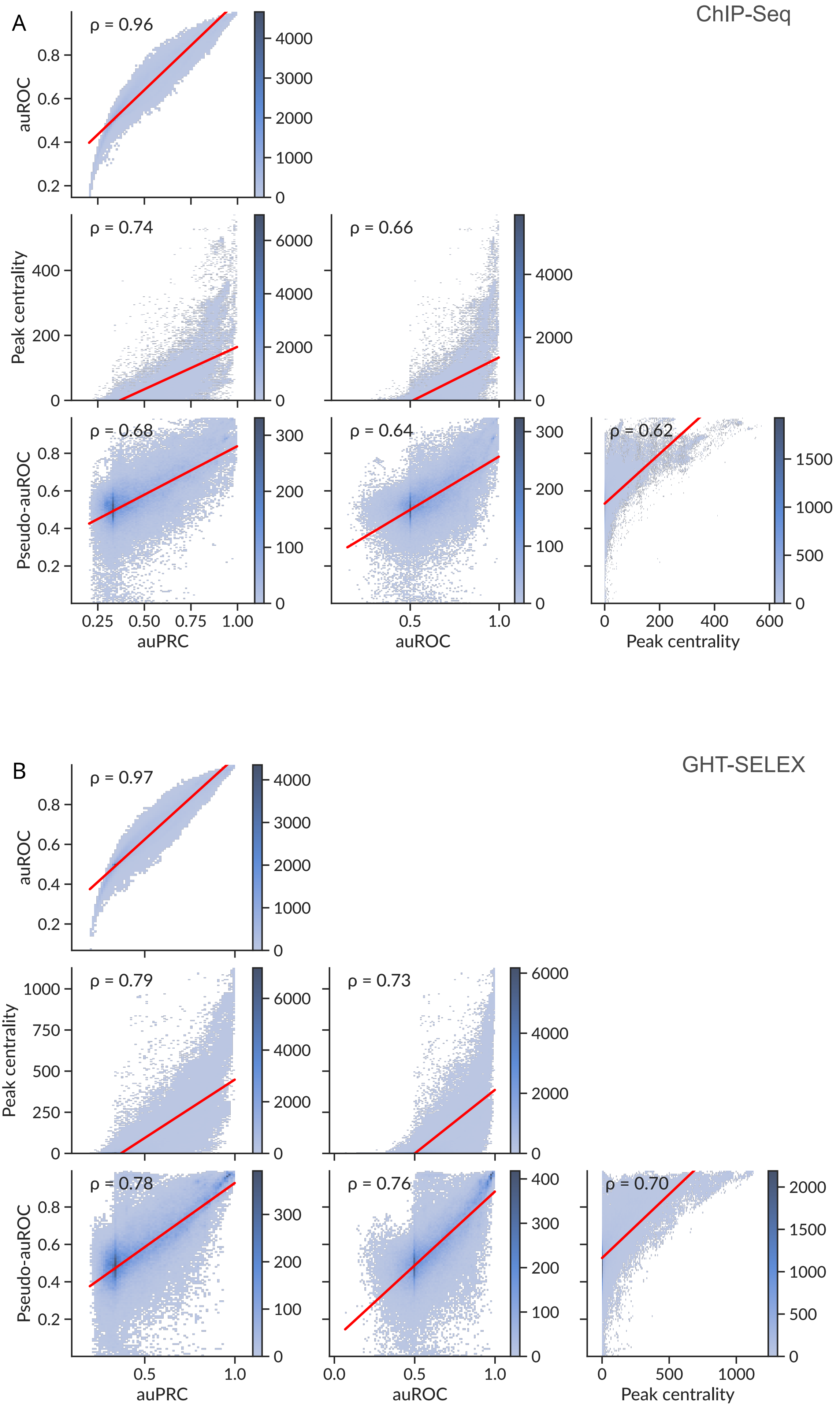


Supplementary Figure SF11
