## Supplementary Figure SF10 for "Cross-platform DNA motif discovery and benchmarking to explore binding specificities of poorly studied human transcription factors"

### Codebook/GRECO-BIT Motif Explorer Website Overview

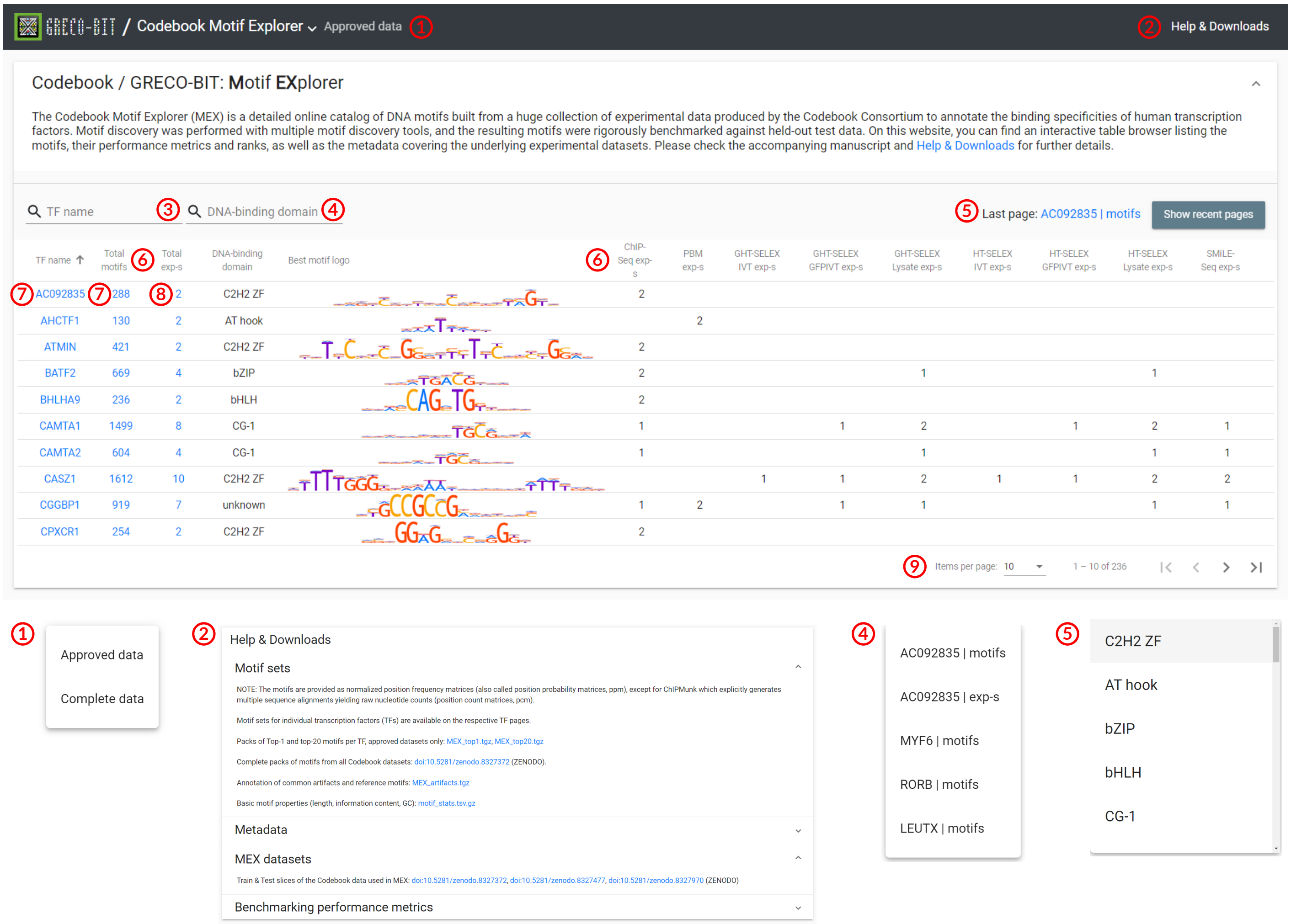


① Swtich between **Approved** and **Complete** sets of motifs and experiments

② Link to the brief **Help** and **Downloads** page

③ Filter the list of TFs and representative motifs by **TF name** or ④ **DNA-binding domain**

⑤ Quick **Page navigation** menu

⑥ Sort the table by clicking on **column headers** (such as the number of experiments a particular type)

⑦ Link to the page listing the **top-ranked motifs** of a particular **TF**

⑧ Link to the page with the **experiment-level metadata**

⑨ Table **pagination** settings

Supplementary Figure SF10

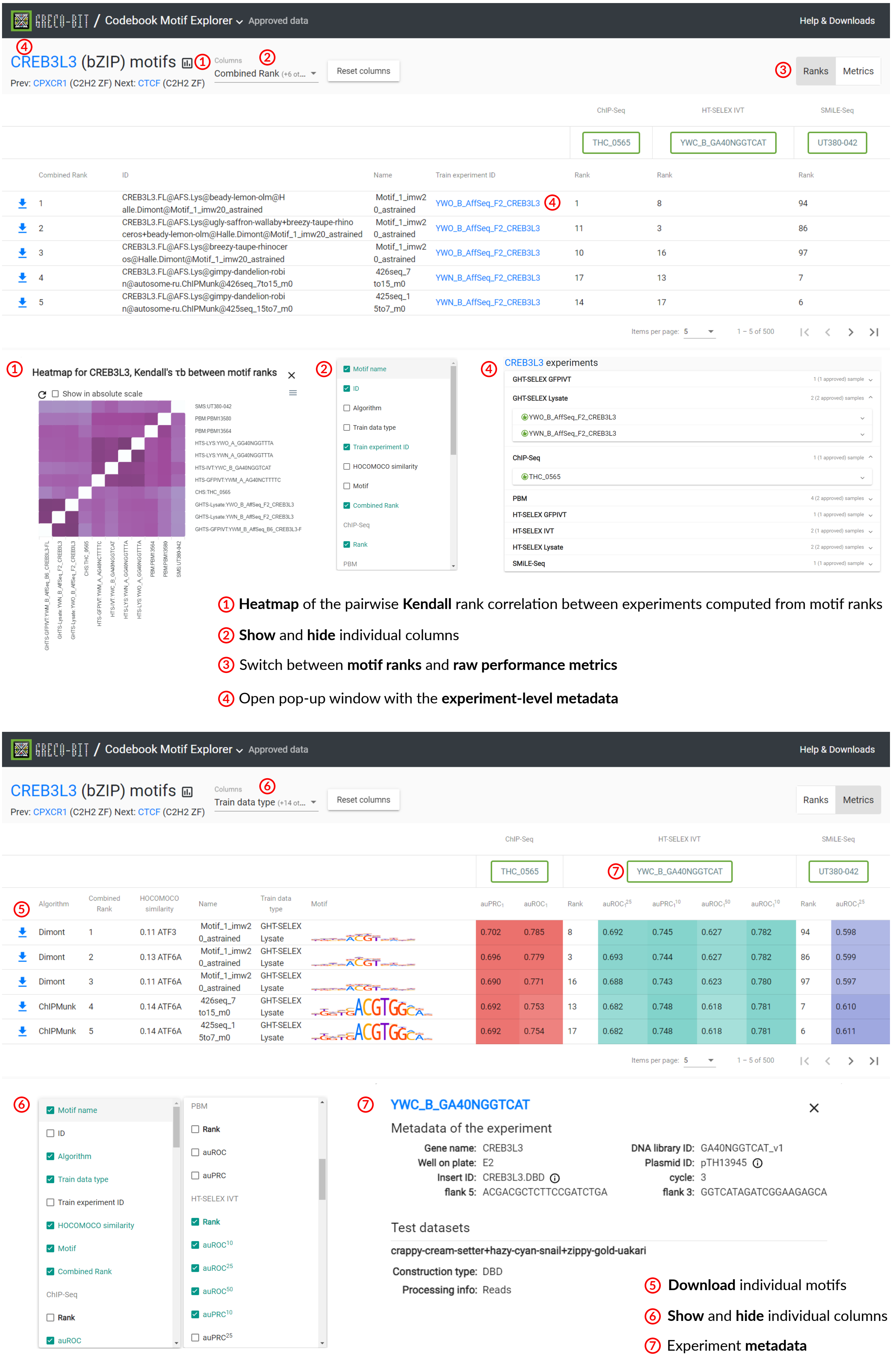


Supplementary Figure SF10 (cont.)
